# Ontogeny and Vulnerabilities of Drug-Tolerant Persisters in HER2+ Breast Cancer

**DOI:** 10.1101/2020.08.28.273029

**Authors:** Che-wei Anderson Chang, Jayu Jen, Shaowen Jiang, Azin Sayad, Arvind Singh Mer, Kevin R. Brown, Allison Nixon, Avantika Dhabaria, Kwan Ho Tang, David Venet, Christos Sotiriou, Jiehue Deng, Kwok-kin Wong, Sylvia Adams, Peter Meyn, Adriana Heguy, Jane Skok, Aristotelis Tsirigos, Beatrix Ueberheide, Jason Moffat, Abhyudai Singh, Benjamin Haibe-Kains, Alireza Khodadadi-Jamayran, Benjamin G. Neel

## Abstract

Resistance to targeted therapies is an important clinical problem in HER2-positive (HER2+) breast cancer. “Drug-tolerant persisters” (DTPs), a sub-population of cancer cells that survive via reversible, non-genetic mechanisms, are implicated in resistance to tyrosine kinase inhibitors (TKIs) in other malignancies, but DTPs following HER2 TKI exposure have not been well characterized. We found that HER2 TKIs evoke DTPs with a luminal-like or a mesenchymal-like transcriptome. Lentiviral barcoding/single cell RNA-sequencing reveal that HER2+ breast cancer cells cycle stochastically through a “pre-DTP” state, characterized by a G_0_-like expression signature and enriched for diapause and/or senescence genes. Trajectory analysis/cell sorting show that pre-DTPs preferentially yield DTPs upon HER2 TKI exposure. Cells with similar transcriptomes are present in HER2+ breast tumors and are associated with poor TKI response. Finally, biochemical experiments indicate that luminal-like DTPs survive via estrogen receptor-dependent induction of *SGK3,*leading to rewiring of the PI3K/AKT/mTORC1 pathway to enable AKT-independent mTORC1 activation.

**STATEMENT OF SIGNIFICANCE:** DTPs are implicated in resistance to TKIs, other targeted therapies, and chemotherapy, but their ontogeny and vulnerabilities remain unclear. We find that HER2 TKI-DTPs emerge from stochastically arising primed cells (“pre-DTPs”) that preferentially engage either of two distinct transcriptional programs upon TKI exposure. Our results provide new insights into DTP ontogeny and identify potential therapeutic vulnerabilities.

## INTRODUCTION

HER2 (a.k.a. ERBB2) drives 15-20% of breast cancers, some of which co-express estrogen receptor (ER). Treatment of non-metastatic HER2+ breast cancer with surgery, chemotherapy, anti-HER2 monoclonal antibodies (trastuzumab + pertuzumab) and, for HER2+/ER+ tumors, anti-estrogens, is a major oncologic success story: 5-year disease-free survival approaches 100% for lymph node (LN)-negative disease and 80-90% (depending on ER status) for patients with local LN involvement (1). By contrast, metastatic HER2+ breast cancer remains incurable (2, 3). Lapatinib, the first clinically approved HER2-directed TKI, inhibits HER2, including antibody-resistant variants, as well as EGFR (4, 5). Combined with chemotherapy, lapatinib improves outcome in patients who progress on trastuzumab/chemotherapy (6). Also, unlike HER2- antibodies, which poorly cross the blood brain barrier (7), lapatinib (and other HER2 TKIs) can target brain metastases. Newer HER2-TKIs, such as neratinib and tucatinib, have more favorable pharmacologic properties and the latter, in particular, is likely to replace lapatinib in metastatic breast cancer regimens (2,8,9).

Multiple mechanisms for lapatinib resistance have been described, including HER3 up- regulation, which increases signaling via HER2/HER3 heterodimers (10–12), activating *PIK3CA* or loss-of-function *PTEN* mutations (13), or increased upstream/parallel pro-survival signaling mediated by FAK1, SRC, PRKACA, or mTORC1 (14–18). Epigenetic mechanisms for lapatinib resistance involving the mixed lineage leukemia (MLL) complex and bromodomain extra terminal domain (BET) family members also have been proposed (19, 20). Although not established by direct experiments, these mechanisms will likely limit the efficacy of other HER2 TKIs as well.

The above events confer stable resistance to lapatinib, but whether resistant cells pre-exist in HER2+ tumors or emerge during therapy remains unclear. Sharma *et al.* described a non-genetic, drug-tolerant cell state in the *EGFR^mut^* non-small cell lung cancer (NSCLC) line PC9, which could be reversed upon drug withdrawal (21). Such cells, which they termed “drug-tolerant persisters” (DTPs), subsequently were observed in other experimental models of targeted therapy, including additional *EGFR*-mutant NSCLC lines (21), *MET*-amplified gastric cancer (22), *BRAF*-mutant melanoma (23–26), *AR-*driven prostate cancer (27, 28), and most recently, after chemotherapy for multiple carcinomas (29, 30) and acute myeloid leukemia (31). Upon continued exposure to EGFR inhibitor, Sharma *et al.* noted that PC9 cells regained the ability to proliferate; they termed such proliferative cells “drug-tolerant expanding persisters” (DTEPs). DTEPs have also been observed in other cell systems (21,26,30–33), and are likely to emerge from the very recently identified “cycling persisters” that comprise a small fraction of the initial DTP population (32). DTPs (and DTEPs) do not appear to be classical “cancer stem cells,” but whether all cancer cells are at any given time equally capable of becoming DTPs remains largely unknown. Also, although epigenetic modulators (e.g., HDAC or KDM5 inhibitors) that prevent development of DTPs in response to EGFR-TKIs have been identified (34), the signaling pathways that DTPs employ to survive TKI treatment are not well understood.

A few studies have identified DTPs in HER2+ breast cancer lines (21,32,35), but they have not been characterized extensively. Here, we provide insights into the ontogeny and potential therapeutic vulnerabilities of HER2 TKI-DTPs. As such cells might comprise a reservoir for the development of stable resistance to HER2-targeted TKIs, our results have potential therapeutic implications.

## RESULTS

### HER2 TKI induce DTPs in some, but not all, HER2-positive breast cancer cell lines

We first asked whether HER2+ breast cancer cells exhibited DTP-like behavior in response to HER2 TKIs. Ten HER2+ breast cancer lines were treated with 2.5 μM lapatinib, a concentration that corresponds to average peak plasma levels in patients (36). Three types of response were observed: (I) 3/10 lines were intrinsically resistant and proliferated in the presence of lapatinib; (II) 2/10 showed a cytostatic response; and (III) in 5/10 lines, most cells died after exposure to lapatinib, but a subpopulation persisted, showing similar behavior to DTPs as defined initially by Sharma *et al*. (Fig. 1A). As noted above, activating *PIK3CA* mutations or *PTEN* deletion can confer lapatinib resistance (13, 37). All Type I and Type II cell lines harbor common “hotspot” activating mutations in *PIK3CA* (H1047R, E545K) or deletion of *PTEN*. Although Type III lines have intact *PTEN* and normal PTEN expression, two feature rare *PIK3CA* variants, encoding K111N (BT474) and C420R (EFM192A) (38). The C420R mutant has increased kinase activity and transforming activity; the pathologic significance of the K111N allele is unclear (39, 40). Apparently, *PIK3CA* mutations can contribute to stable lapatinib resistance, yet not all such mutations are sufficient to confer resistance.

**Figure 1.**
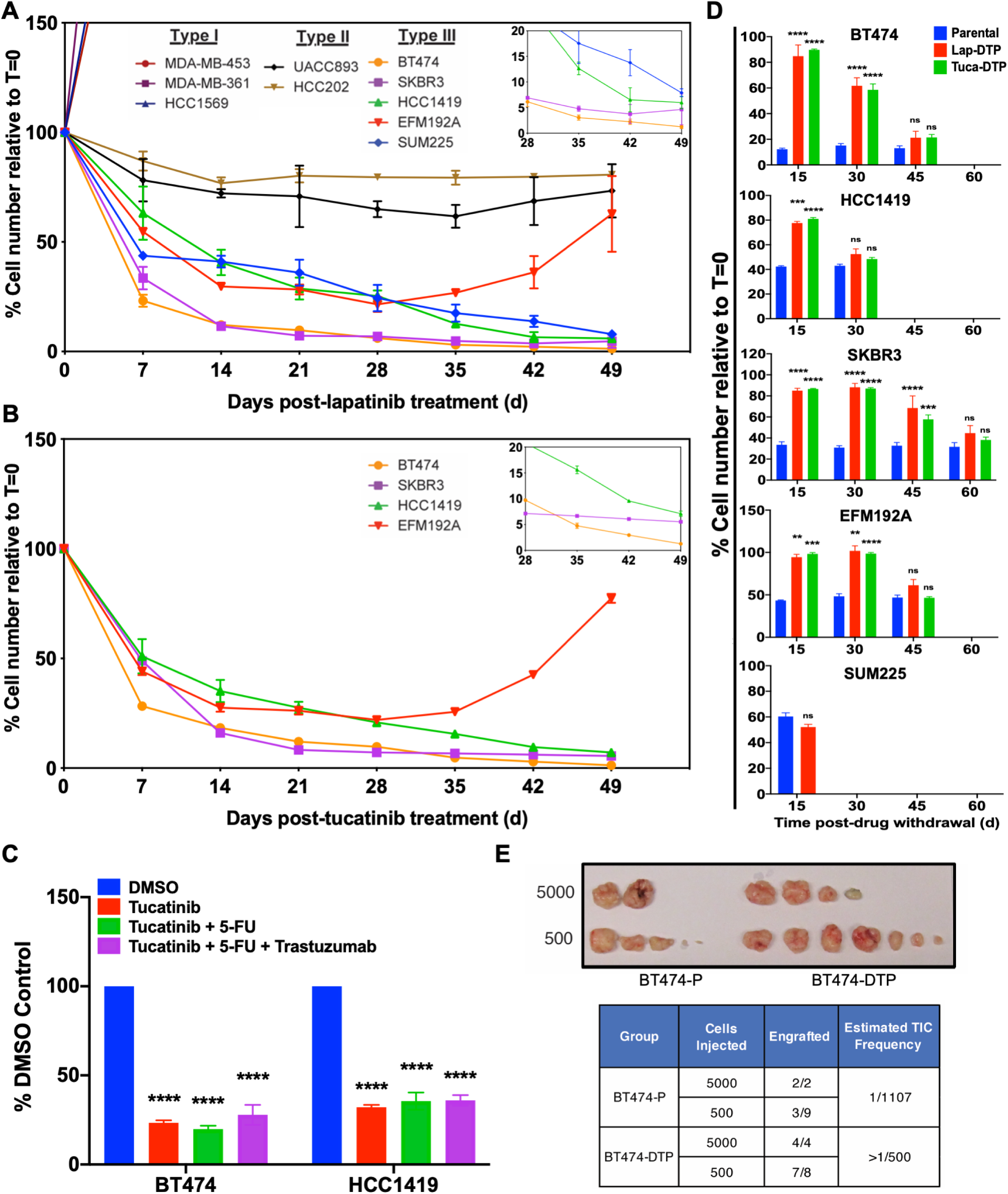
Some HER2+ breast cancer cell lines give rise to HER2 TKI drug-tolerant persisters (DTP). **A** and **B,** HER2+ breast cancer cell lines were treated with 2.5 μM lapatinib (**A**) or with 1.2 μM tucatinib (**B**), counted at the indicated times, and the percentage of the initial cell number was determined. Mean ± SEM from three independent experiments is displayed shown. The inset shows a magnified view of the residual cells from Days 28-49. **C,** BT474 and HCC1419 cells were treated with the indicated agents for 14 days and percentage survival was quantified. **D,** Cells from the indicated lines were cultured in lapatinib or tucatinib (as indicated) for 14 days to generate DTPs. Then drug was withdrawn, and cells were allowed to resume proliferation. At the indicated times, cells were re-challenged with the same drug. Parental cells were used as a control at each time point. Cell survival was assessed by counting viable cells after 7-days of TKI treatment and normalized to the viable cell number at Day 0 (t=0). Mean ± SEM from three independent experiments is displayed. Statistical significance was assessed by two-tailed t-test for SUM225. All other cell lines were assessed by two-way ANOVA with Bonferroni post-hoc analysis (ns, p>0.05, *p<0.05 **p<0.01, ***p<0.001, ****p<0.0001). **E,** Number of tumors detected and estimated tumor-initiating cell (TIC) frequency of BT474 parental cells and lapatinib-DTPs after injection into the mammary fat pad of NSG mice. The top image shows tumors 5 months post-injection.

Unlike the behavior of the lines tested by Sharma *et al.* (21), only EFM192A cells entered a DTEP-like state, which was evident after 30 days of continuous drug treatment. The other Type III lines (BT474, SKBR3, HCC1419, SUM225) remained quiescent for ∼50 days (Fig. 1A), a period during which Sharma *et al.* observed DTEPs emerging from EGFR inhibitor-treated PC9 cells. Type III lines also gave rise to DTPs in response to the next-generation HER2 TKI tucatinib at concentrations corresponding to steady state C_max_ levels (1.2 μM) in patients (Fig. 1B). HER2 TKIs are rarely administered as single agents to HER2+ breast cancer patients, so we tested clinically relevant HER2 TKI combinations, including tucatinib, tucatinib + fluorouracil (5-FU), tucatinib + 5FU + trastuzumab. These combinations yielded similar number of residual cells, suggesting that HER2 TKI-DTPs are cross-resistant to conventional HER2 treatment regimens (Fig. 1C).

Upon drug withdrawal, DTPs yield progeny that regain drug sensitivity (21). As expected, “HER2 TKI-DTPs” resumed proliferation after lapatinib or tucatinib withdrawal, demonstrating that they were not permanently growth arrested. At various times after drug withdrawal, we rechallenged these cells with lapatinib or tucatinib. Consistent with their classification as DTPs, all Type III cells that survived initial exposure to lapatinib or tucatinib yielded HER2 TKI-sensitive progeny upon drug withdrawal (Fig. 1D). Individual cell lines differed in the time required to regain drug-sensitivity, but this interval was highly reproducible across several experiments. Furthermore, BT474-derived lapatinib-DTPs were at least as tumorigenic (if not more so) as parental BT474 cells, as assessed by a limiting dilution assay in NOD.Cg-Prkdc^scid^ Il2rγ^tm1Wjl^/SzJ (NOD scid gamma, NSG) mice (Fig. 1E). Hence, not only do these cells have the potential to restart proliferation following HER2 TKI withdrawal, they also can seed new tumors.

### HER2 TKI-DTPs display two distinct transcriptional profiles

We next compared the transcriptomes of HER2 TKI-DTPs and parental cells by bulk RNA sequencing (RNA-seq). Unsupervised hierarchical clustering of DTPs remaining after lapatinib treatment (lapatinib-DTPs) segregated the samples into two subgroups, each of which differentially expressed distinct sets of genes compared with their parental counterparts (Fig. 2A; Supplementary Tables 1-2). Gene Set Enrichment Analysis (GSEA) using a compendium of pathway gene sets compiled by the Bader laboratory (41) revealed that lapatinib-DTPs in one cluster differentially activated a gene set annotated as “Hallmark Epithelial Mesenchymal Transition”; we refer to these cells as “mesenchymal-like” DTPs (Fig. 2B; Supplementary Fig. S1A). DTPs from the other cluster activated “Hallmark Estrogen Response Early” genes and hereafter are termed “luminal-like” (Fig. 2B; Supplementary Fig. S1B). The differentially expressed genes (DEGs) in DTPs and parental cells from the two subgroups enriched for distinct sets of transcription factor binding sites by Chip Enrichment Analysis (ChEA): mesenchymal-like DTPs were enriched for SMAD4 and SOX2 sites, whereas DEGs in luminal-like DTPs showed enrichment for ER (ESR1) sites (Supplementary Fig. S1C-F). Tucatinib treatment also evoked DTPs with either luminal-like or mesenchymal-like transcriptomes (Fig. 2C, Supplementary Fig. S1G). Supervised analyses revealed markedly similar transcriptomic changes in DTPs induced by tucatinib or lapatinib, although the next generation TKI more strongly induced or repressed many individual DEGs (Fig. 2D).

**Figure 2.**
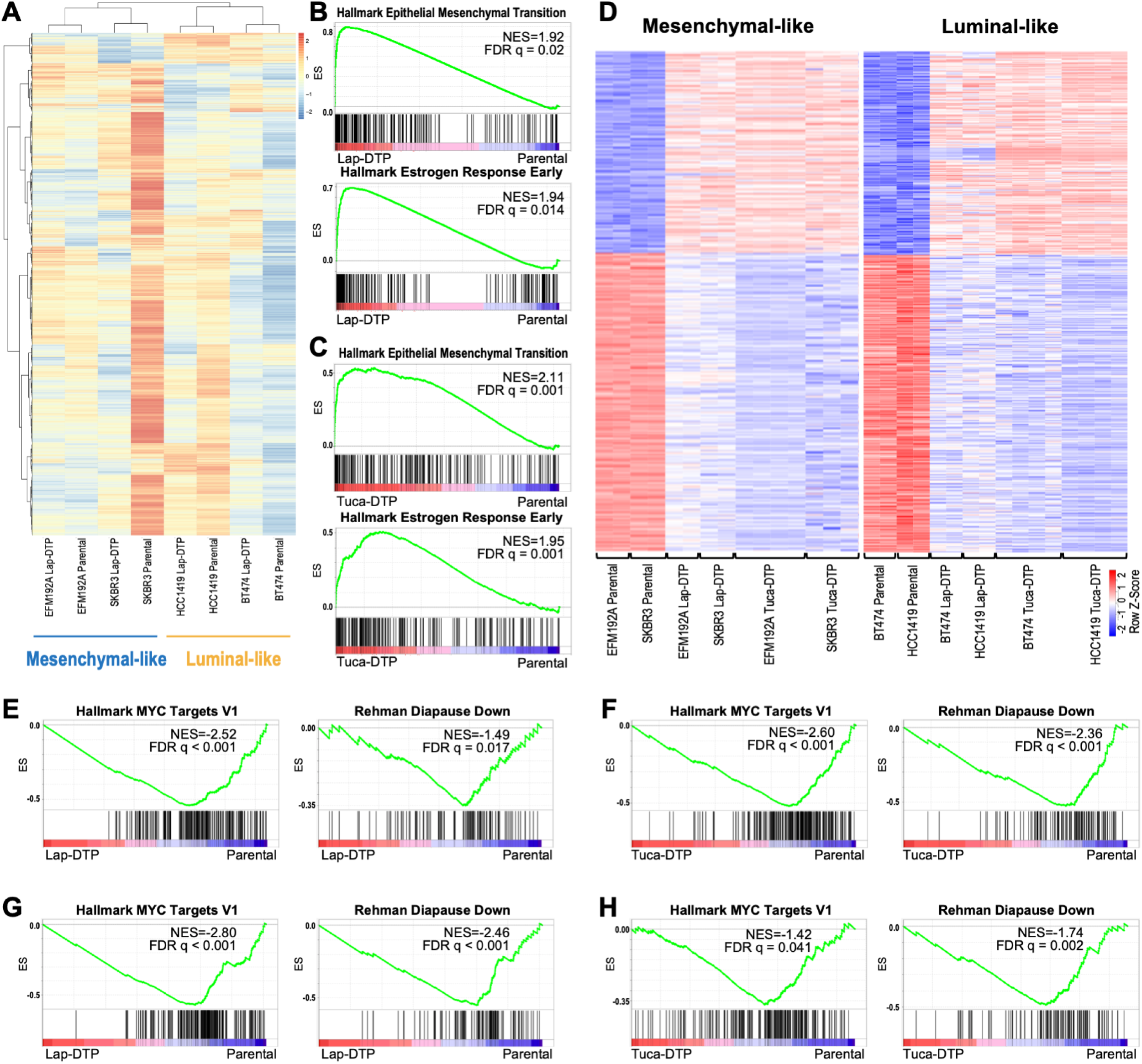
HER2+ breast cancer cells elicit distinct transcriptional programs after HER2 TKI treatment. **A**, Heatmap shows unsupervised clustering of parental and lapatinib-DTP samples. Scale represents the z-score. **B** and **C,** GSEA shows enrichment of “Hallmark Epithelial Mesenchymal Transition” genes in mesenchymal-like DTPs (top) and “Hallmark Estrogen Responses Early” in luminal-like DTPs (bottom) evoked by lapatinib (**B**) or tucatinib (**C**). **D,** Heatmaps show supervised clustering of parental, lapatinib-DTPs and tucatinib-DTPs with mesenchymal-DTP DEGs (left) or luminal DTP DEGs (right). Scale represents the z-score. **E**-**H**, GSEA shows enrichment of the indicated gene sets in lapatinib-induced (E) and tucatinib-induced (F) mesenchymal-like DTPs, and lapatinib-(G) and tucatinib-induced (H) luminal-like DTPs.

While this manuscript was in revision, several studies reported that treatment of multiple types of carcinomas or acute myeloid leukemia with chemotherapy induce DTPs that down-regulate MYC target genes and induce embryonic diapause and/or senescence-like transcriptional programs (29–31). Only some of these gene sets are represented in the Bader collection (see above). Notably, supervised analysis revealed that expression of “Hallmark MYC Targets” and “Rehman Diapause Down” (29) genes decreased significantly in lapatinib- and tucatinib-evoked mesenchymal-like and luminal-like DTPs compared with their respective parental cells (Fig. 2E-H). Other signatures from these recent reports, including chemotherapy-induced stress genes (“Duy CISG”) (31) and senescence genes (“Fridman Senescence”), were enriched in mesenchymal-like (but not luminal-like) lapatinib- and tucatinib-DTPs (Supplementary Fig. S1H-K).

### Lapatinib-DTPs are organized stochastically

To clarify their ontogeny, we asked whether lapatinib-DTPs belong to a pre-existing cellular hierarchy (e.g., a “cancer stem cell” model) or arise stochastically. We transduced a lentiviral barcode library into BT474 cells at a low multiplicity of infection (MOI=0.1) to ensure that each cell received only one barcode; this approach enabled tracking of up to 100,000 unique clones (Supplementary Fig. S2A and Methods). Infected cells were permitted to expand for approximately six weeks, so that each barcode was represented abundantly in the overall cell population. Approximately 12% of BT474 cells survive 14 days of lapatinib treatment (Fig. 1A). If HER2+ breast cancer lines maintain a pre-existing, fixed hierarchy of DTPs and non-DTPs, a similar percentage of barcodes should be retrieved after 14-day lapatinib treatment, compared to the number of barcodes in the initial population (t=0). By contrast, if lapatinib-DTPs arise stochastically, substantially >12% of barcodes should be retrieved (Supplementary Fig. S2A).

In two independent experiments, we retrieved 62% and 60% of barcodes from transduced BT474 cells, respectively (Supplementary Fig. 2B, left two panels). We noted a consistent reduction (∼25%) in barcode representation in untreated control (UT) cells cultured for 14 days without lapatinib in both experiments. These “missing barcodes” were poorly represented at t=0, and presumably decreased to undetectable levels after 14 days in culture. Others have also reported stochastic loss of barcodes upon cell passaging (42). Nevertheless, the retention of most barcodes in the starting population after 14 days of lapatinib indicates that nearly all BT474 cells can give rise to lapatinib-DTPs. Similar results were obtained in analogous single experiments on HCC1419, SKBR3, and EFM192A cells (Supplementary Fig. 2B, right three panels). These data support a stochastic model of DTP ontogeny and suggest that over a 6-week period (or less), essentially every HER2+ cell or its progeny has the capacity to transit into the drug-tolerant state.

### Parental BT474 cells occupy states with different predilections to become DTPs

Although the barcoding experiments established that all cells can (over a 6-week period) occupy a state that can become a DTP upon HER2 TKI exposure, fluctuation testing (43) indicated that at any given time, only some cells (“pre-DTPs”) are primed to become DTPs. Interestingly, this analysis also suggests that, depending on the specific cell line, the pre-DTP state is heritable for ∼2-7 generations (Supplementary Fig. 2C).

To investigate further the dynamics of DTP generation, we performed single-cell RNA sequencing (scRNA-seq) on untreated (UT), 6-hour lapatinib-treated (6h), and 14-day lapatinib-treated BT474 cells (DTPs) and analyzed the data with iCellR (44). The 6-hour timepoint was chosen because lapatinib-treated BT474 cells began to die at this time. In supervised analyses using the DEGs in BT474-DTPs (versus parental cells) from bulk RNA-seq (“BT474-DTP DEG”; Supplementary Table 3), we observed that most DEGs that are increased in BT474 DTPs (BT474-DTP Up DEGs) were induced, while most DEGs that are decreased in BT474 DTPs (BT474-DTP-Down DEGs) were down-regulated progressively upon lapatinib treatment (Fig. 3A). However, closer examination revealed bi-modal expression of BT474-DTP Down DEGs in untreated BT474 cells (compare violin plots of UT cells using BT474-DTP Up and BT474-DTP Down DEGs, respectively). This finding, along with the barcoding and fluctuation results described above, argues that only select cells exist in a state (“pre-DTPs”) conducive to DTP generation.

**Figure 3.**
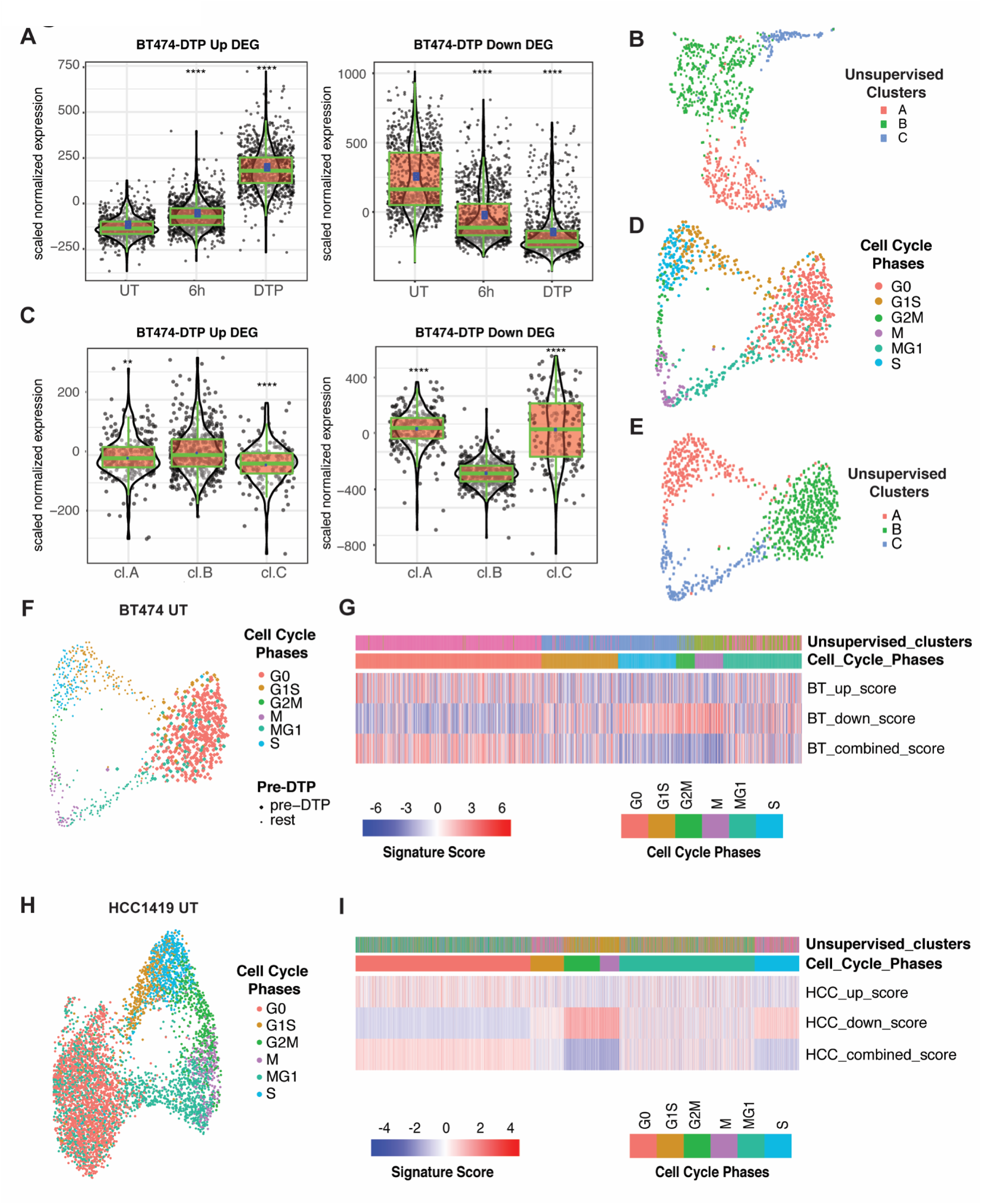
A fraction of randomly growing HER2-luminal cells occupies a pre-DTP state characterized by a G_0_-like signature. **A**, Aggregate expression of BT474 lapatinib-DTP Up and Down DEGs in single cells from untreated (UT), 6-hour lapatinib-treated (6h) and DTP (14-day lapatinib) samples. ****P<0.0001, unpaired t-test compared to UT. **B,** UMAP projection of scRNA-Seq of untreated BT474 cells. Cells are colored by their unsupervised clusters. **C,** Aggregate expression of BT474 lapatinib-DTP Up and Down DEGs, as indicated, in each unsupervised cluster of untreated BT474 cells from **B**. **P<0.01, ****P<0.0001, unpaired t-test compared to cluster B. **D,** UMAP projection of untreated BT474 cells using cell cycle signature genes as defined by Xue *et al.* with cells colored by cell cycle stage (see Methods for details). **E,** UMAP projection showing untreated BT474 cells clustered by the Xue *et al.* cell cycle genes (as in **D**), but with cells colored according to their unsupervised clusters (as determined in **B**). **F,** Same projection as in **D**, but with pre-DTPs (based on their BT474-DTP combined DEG score) shown as larger circles compared with other cells. **G,** Heatmap displaying BT474-DTP Up DEG score, BT474-DTP Down DEG score and BT474-DTP combined DEG score (up DEG score minus down DEG score), in each untreated BT474 cells. The upper panels show the unsupervised clusters from **B** and their cell cycle phase, determined by expression of the Xue *et al*. cell cycle genes. **H,** UMAP projection of supervised clustering of untreated HCC1419 cells by Xue *et al.* cell cycle genes. Cells are colored by cell cycle status. **I,** Heatmap displaying HCC1419-DTP Up DEG score, HCC1419-DTP Down DEG score, and HCC1419-DTP combined DEG score (up minus down DEG score) in single cells from HCC1419. Upper panels display the unsupervised cluster to which each untreated HCC1419 cell belongs and its inferred cell cycle phase.

To explore this intriguing possibility, we performed additional analyses on untreated (UT) BT474 cells. Unsupervised clustering identified three major clusters (Fig. 3B), of which cluster B alone displayed significant enrichment for both up- and down-regulated BT474-DTP DEGs compared with the other two (Fig. 3C). Gene ontology (GO) enrichment analysis (using Enrichr) showed that the unsupervised clusters were distinguished by genes associated with distinct cell cycle phases (Supplementary Fig S3A; Supplementary Tables 4 and 5). Cluster A showed enrichment for genes involved in the G1/S transition, whereas cluster C was enriched for G2/M genes. Additional supervised analyses using signatures that can distinguish multiple stages of the cell cycle, including G_0_ (45), revealed strong overlap between cells in the putative “pre-DTP” cluster (cluster B) and G_0_ cells (Figs. 3D-F). Moreover, most G_0_ cells and a few cells annotated as “G1S or MG1” differentially expressed BT474-DTP Up and Down DEGs as well as luminal-like DTP Up and Down DEGs (Fig. 3G; Supplementary Figs. S3B-E). The other luminal-like cell line, HCC1419, also contained a cluster with a G_0_ signature, increased expression of HCC1419-DTP Up and luminal-like Up DEGs and decreased expression of HCC1419-DTP Down and luminal-like Down DEGs (Figs. 3H and 3I; Supplementary Figs. S3G-L; Supplementary Table 6). To remove proliferation-related genes that might be differentially expressed in DTPs (which are non-proliferative), we removed overlapping DEGs in luminal-like and mesenchymal-like DTPs (Supplementary Fig. S1C). The resultant “Mesenchymal DTP Unique DEGs” (Supplementary Table 7) and “Luminal DTP Unique DEGs” (Supplementary Table 8) also were enriched in a sub-population of G_0_-like cells from all untreated mesenchymal-like and luminal-like lines examined (Fig. 4A).

**Figure 4.**
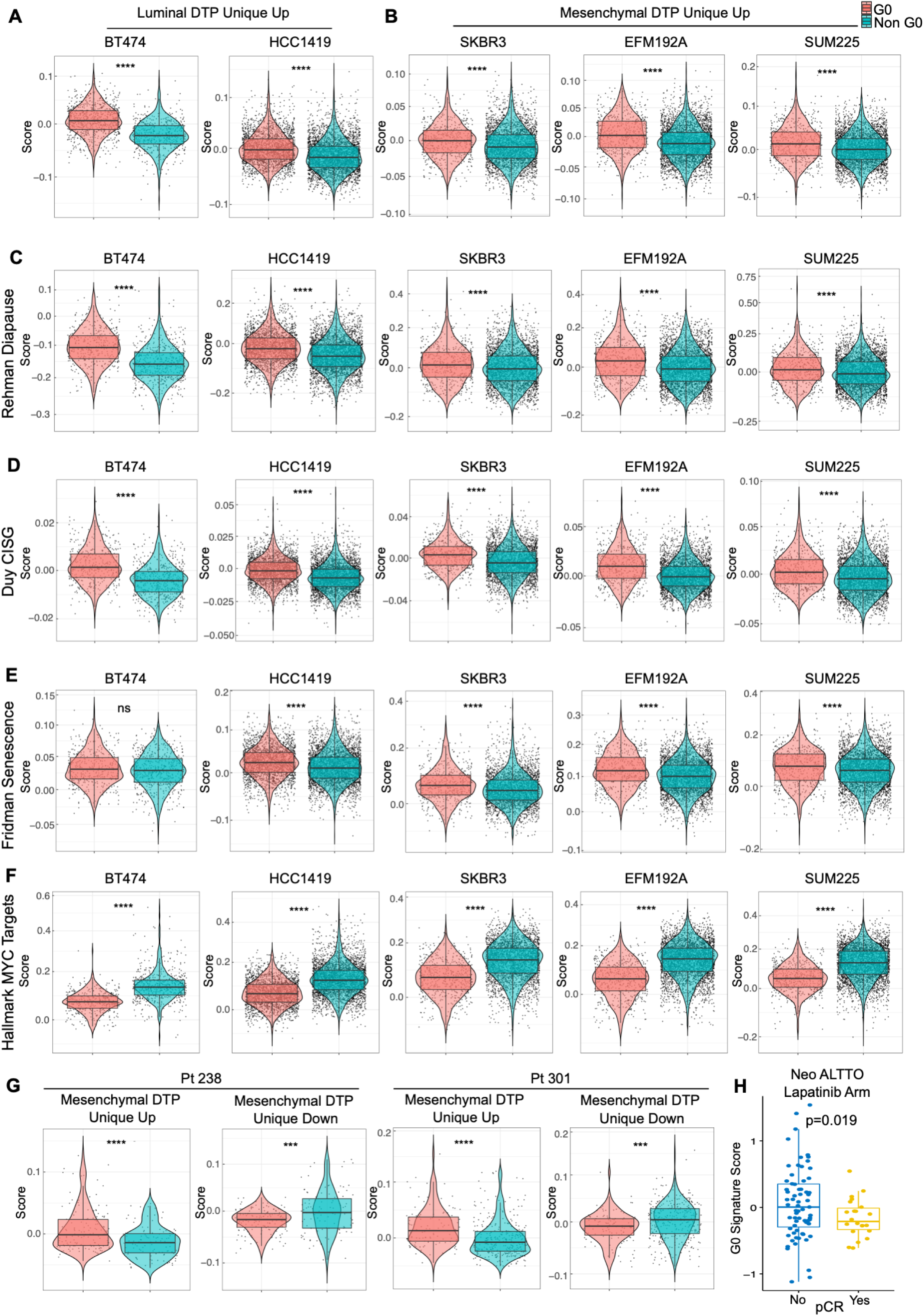
G_0_-like, pre-DTP cells in HER2+ breast cancer cell lines and patients enrich for gene signatures associated with other DTPs and anti-correlate with pathological response rate (pCR) to lapatinib. A and **B,** Cells with G_0_ signature in luminal-like HER2+ breast cancer lines, BT474 and HCC1419, are enriched for Luminal DTP Unique Up genes (**A**), while cells with G_0_ signature in mesenchymal-like HER2+ breast cancer lines, SKBR3, EFM192A and SUM225, are enriched for Mesenchymal DTP Unique Up genes (**B**). G_0_-like cells were identified by using cell cycle signatures from Xue *et al.* C-**F,** G_0_ cells in all HER2+ breast cancer cell lines are enriched for Rehman Diapause Up (29) (**C**), Duy CISG Up (31) (**D**), and Fridman Senescence Up (31, 46) genes (**E**) and depleted for Hallmark MYC Target genes (**F**). **G,** G_0_ cells in two primary HER2+/ER-breast cancer patients are enriched for Mesenchymal DTP Unique Up genes. **H**, Distribution of G_0_ signature scores calculated from Oki *et al.* (91) in patients with or without pCR in the neoadjuvant lapatinib arm of the NeoALTTO trial, analyzed by two-sided Student’s t-test.

Collectively, these data show that randomly proliferating HER2+ breast cancer cell lines stochastically maintain a population of cells characterized by a G_0_-like signature and expression of a subset of genes differentially expressed in DTPs, even though these cells have never been exposed to HER2 TKIs. Intriguingly, compared with non-G_0_ cells, G_0_-like cells from multiple HER2+ breast cancer lines also were enriched for diapause, chemotherapy-induced stress genes (CISG) (31), and senescence gene sets (31, 46) and were anti-correlated with “Hallmark MYC Targets” (29–31) (Fig. 4C-F). Furthermore, scRNA-seq data from HER2+/ER-tumors from two treatment-naïve patients (Pt 238 and Pt 301) revealed the presence of G_0_-like cells that were enriched for Mesenchymal DTP Unique Up and depleted for Mesenchymal-DTP Unique Down DEGs compared with non-G_0_ cells (Fig. 4G). In general, compared with non-G_0_ cells, G_0_-like cells from these tumors also were enriched for diapause, CISG, and senescence genes and showed down-regulation of “Hallmark MYC Targets” genes (Supplementary Fig. S4A and S4B). Importantly, analysis of bulk RNA-seq data from HER2+ breast cancer patients in the NeoALTTO trial who did not attain pathological complete response (pCR) after neo-adjuvant lapatinib treatment had higher G_0_ signature scores compared with those achieving pCR (Fig. 4H). These results indicate that G_0_-like cells with some transcriptional features of DTPs pre-exist in tumors from untreated HER2+ breast cancer patients and are negatively associated with HER2 TKI response, comporting with the possibility that these cells become DTPs upon drug exposure. Remarkably, cells with similar transcriptional properties also were detectable in tumors from untreated triple negative (TN) breast cancer patients, raising the possibility that a G_0_-like pre-DTP state might exist in other breast cancer subtypes (Supplementary Fig. S4C and S4D).

Seeking functional evidence for the proposed G_0_-like “pre-DTP” state, we searched for cell surface markers differentially expressed in cluster B (pre-DTPs) compared with the other two clusters. Twenty DEGs were up-regulated in cluster B and in BT474-DTPs, including several encoding surface proteins: *NPY1R*, *MAN1A1, ABCC5*, *CLDN8, PSCA*, and *PIK3IP1* (Fig. 5A, Supplementary Figs. S5A and S5B). Using an anti-NPY1R antibody, we purified BT474 cells with low, medium, and high surface expression of NPY1R by fluorescence-activated cell sorting (FACS) (Supplementary Fig. S5C). Remarkably, untreated BT474 cells with successively higher NPY1R expression also showed successively increased resistance to lapatinib (Fig. 5B; Supplementary Fig. S5D). Consistent with the stochastic origin of lapatinib-DTPs revealed by the barcoding experiments, cells purified based on different levels of surface NPY1R recapitulated heterogeneous NPY1R expression after 14 days in culture (Supplementary Fig. S5E). These fractions also proliferated similarly post-sort (Supplementary Fig. S5F), ruling out the possibility that contaminating NPY1R^hi^ cells had overtaken the initially NPY1R^lo^ cell population or *vice versa*. We also tested another surface marker preferentially expressed in DTPs and putative pre-DTPs, ABCC5. ABCC5^hi^ BT474 cells were substantially enriched for NPY1R^hi^ cells compared with the ABCC5^lo^ fraction, indicating co-expression of both markers in a subset of BT474 cells (Supplementary Fig. S5G). Compared with the ABCC5^lo^ fraction, ABCC5^hi^ cells also had increased ability to generate DTPs in response to lapatinib or tucatinib treatment (Supplementary Fig. S5H-J). Bulk RNA-seq analysis revealed ∼500 DEGs in FACS-enriched NPY1R^hi^ versus NPY1R^lo^ cells (Supplementary Fig. S6A; Supplementary Table 9), and NPY1R^hi^ DEGs were enriched significantly for BT474-DTP DEGs (Fig. 5C).

**Figure 5.**
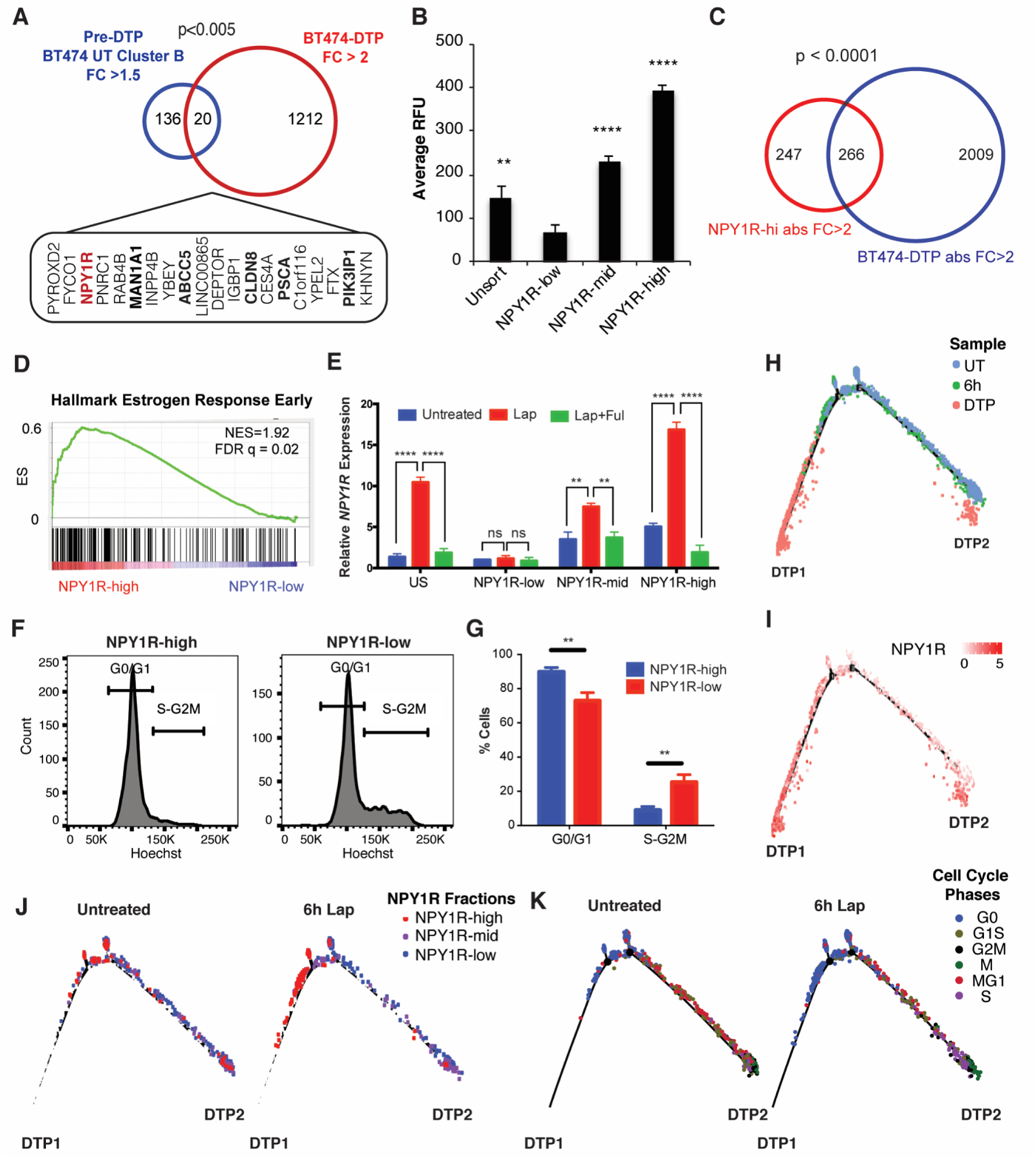
Prospective purification of pre-DTPs. **A**, Overlap of untreated BT474 cluster B DEGs with fold change (FC) >1.5 and BT474 lapatinib-DTP DEGs with FC > 2. The 20 overlapping genes and the p-values from Fisher’s exact test are displayed. Genes encoding surface proteins are shown in bold. **B,** FACS-isolated NPY1R^hi^, NPY1R^mid^, and NPY1R^lo^ cells were treated with lapatinib for 14 days. Average RFU indicates the Alamar Blue readings at the experimental endpoint. Mean ± SEM for three independent experiments is displayed. Significance was evaluated by two-tailed t-test (**p<0.01, ****p<0.0001). **C,** Overlap of NPY1R and BT474 lapatinib-DTP DEGs with directionality of the DEGs matched in the analysis and p-value from Fisher’s exact test is displayed. **D,** GSEA shows enrichment for “Hallmark Estrogen Response Early” genes in NPY1R^hi^ versus NPY1R^lo^ cells. **E,** qRT-PCR for *NPY1R* in FACS-isolated NPY1R^hi^, NPY1R^mid^, and NPY1R^lo^ cells that were either left untreated, treated with lapatinib for 6 hours (Lap), or treated with lapatinib and fulvestrant for 6 hours (Lap + Ful). Relative *NPY1R* expression represents the 2^-ΔΔCT^ relative to the untreated NPY1R^lo^ sample. The mean ± SEM of three independent experiments is displayed (ns, p>0.05, **p<0.01, ****p<0.0001, two-tailed t-test). **F,** BT474 cells were stained with Hoechst 33342 and for surface NPY1R; the NPY1R^hi^ and NPY1R^lo^ populations are displayed. **G,** Quantification of cells in G_0_/G_1_ and S-G_2_M from panel **F** (**p<0.01, two-tailed t-test). **H** and **I,** Pseudotime analysis of cells from untreated BT474 cells, 6h lap-treated cells, and DTP samples with cells colored by sample condition (**H**) or *NPY1R* expression (**I**). **J,** Cells from BT474 untreated and 6h Lap sample were isolated (computationally), and their respective positions on the pseudotime trajectory are displayed. Cells are colored by their relative *NPY1R* expression level. **K,** Cells from untreated and 6h lapatinib-treated BT474 cells were isolated (computationally), and their respective positions on the pseudotime trajectory are displayed. Cells are colored by their cell cycle phase.

As noted above, lapatinib-DTPs from BT474 cells displayed enrichment for the estrogen receptor (ER)-driven transcriptome (Fig. 2B; Supplementary Fig. 1B). Notably, “Hallmark of Estrogen Response Early” genes were the most differentially enriched gene set in NPY1R^hi^, compared with NPY1R^lo^, cells (Fig. 5D; Supplementary Fig. S6B). Multiple ER target genes were expressed at higher levels in NPY1R^hi^ than in NPY1R^lo^ cells, suggesting that these cells have basally higher ER signaling activity prior to lapatinib treatment (Supplementary Fig. S6C). Moreover, TCGA data show that *NPY1R*^hi^ tumors express higher levels of *ESR1* and ER targets (*PGR, GREB1, STC2*) than those with low *NPY1R* expression (Supplementary Fig. S6D). Lapatinib treatment induced *NPY1R* in NPY1R^hi^ cells to even higher levels than in NPY1R^mid^ and NPY1R^lo^ cells (Fig. 5E). NPY1R^hi^ cells also were depleted for genes involved in mitosis or G2/M checkpoint compared with NPY1R^lo^ cells (Supplementary Fig. S6B). Co-staining with NPY1R antibody and Hoechst33342 confirmed that NPY1R^hi^ cells were mainly in G0/G1 whereas NPY1R^lo^ cells distributed across all cell cycle phases (Figs. 5F and 5G). Taken together, these data indicate that NPY1R expression marks G0-like, pre-DTP cells that are primed to become DTPs upon HER2 TKI exposure.

Using scRNA-seq data from the UT, 6h, and DTP samples, we performed “pseudotime analysis” to infer the trajectory by which parental, or 6h-treated, BT474 cells become lapatinib-DTPs. We found that BT474-DTPs separated into two states: nearly all DTPs (>90%) were in “DTP state 1” (DTP1), while a much smaller number occupied “DTP state 2” (DTP2) (Fig. 5H). Cells progressively up-regulated *NPY1R* as they progressed from UT to 6h-treated to DTP cells (Fig. 5I). NPY1R^hi^ cells mainly followed the DTP1 route, whereas cells with lower NPY1R levels favored the DTP2 route (Fig. 5J). Comporting with our finding of a G_0_-like pre-DTP state, BT474 and HCC1419 cells annotated as G_0_ favor the main (DTP1) route in the pseudotime analysis (Fig. 5K; Supplementary Figs. S6E and S6F). Direct comparison of the levels of genes preferentially expressed in the pre-DTP/G_0_-like cluster in UT BT474 cells after 6 hours of lapatinib treatment and in the DTP state, however, reveal different patterns of behavior. For example, expression of some cluster B/G_0_ genes decrease as cells transit to the DTP state, some change little if at all, whereas others are induced (Supplementary Tables 10 and 11). These findings suggest that some genes that appear to be “induced” by lapatinib treatment are actually “selected” by virtue of their pre-existing expression in G_0-_like pre-DTPs, whereas others are primed for induction upon HER2-TKI exposure. Taken together, our data establish cells transiting through G_0_ (or conceivably just entering or leaving that state) can proceed preferentially to the DTP state upon TKI treatment and give rise to the vast majority (DTP1 in Fig. 5) of DTPs (see Discussion).

### Lapatinib-DTPs activate mTORC1 via a PI3K-dependent, AKT-independent pathway

RNA-seq analysis suggested that when HER2 signaling is abrogated by TKI exposure, luminal-like DTPs survive via ESR1 (ERα)-driven signaling. Consistent with this hypothesis, BT474 and HCC1419 cells were killed more effectively by lapatinib plus fulvestrant (Lap + Ful, Figs. 6A and 6B). Almost all BT474 DTPs were eradicated upon combination treatment, as evident by the lack of cell re-growth in Lap + Ful combination compared with lapatinib alone (Fig. 6C). The response to Lap + Ful correlated with basal and induced *ESR1* levels (Fig. 6D). Collectively, these data show that the distinct transcriptional programs evoked in luminal-like and mesenchymal-like DTPs confer different therapeutic vulnerabilities.

**Figure 6.**
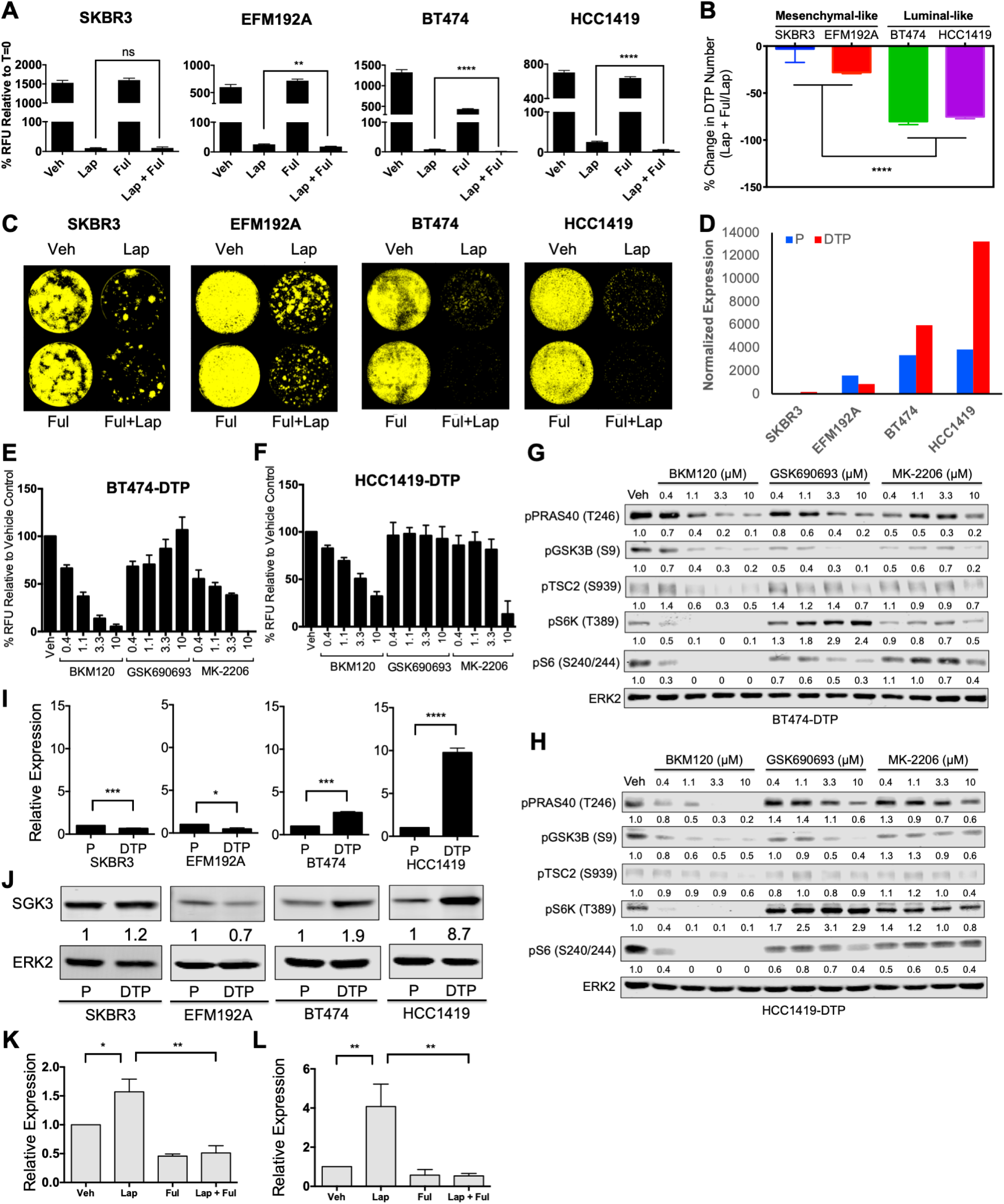
Luminal-like lapatinib-DTPs are more sensitive to PI3K than AKT inhibition and transcriptionally up-regulate *SGK3* via estrogen receptor. **A**, Alamar Blue readings of SKBR3, EFM192A, BT474, and HCC1419 cells treated as indicated for 7 days. Relative fluorescence units (RFU) were normalized to t=0. **B,** Changes in DTP numbers following fulvestrant plus lapatinib treatment compared with lapatinib treatment alone; Mean ± SEM of three independent experiments is displayed. Significance was assessed by two-tailed t-test (****p<0.0001). **C**, Incucyte images of SKBR3, EFM192A, BT474, and HCC1419 cells treated as indicated for 14 days and then cultured drug-free for 14 days to assess regrowth. Representative images from three independent experiments are displayed. **D,** DESeq2 normalized expression of *ESR1* is displayed for parental cells and lapatinib-DTPs of each cell line. **E** and **F,** Lapatinib-DTPs from BT474 (**E**) or HCC1419 (**F**) cells were treated with increasing doses of BKM120 (pan-PI3K inhibitor), GSK690693 (AKT catalytic inhibitor), or MK-2206 (AKT allosteric inhibitor) for 96 hours, and cell number was determined by Alamar Blue viability assay. Mean ± SEM from three independent experiments is shown. **G** and **H,** Differential effect of PI3K and AKT inhibitors on PI3K pathway components. Cells were treated with the indicated inhibitors or vehicle (DMSO) for one hour before lysis. Whole cell lysates were resolved by SDS-PAGE and immunoblotted with the indicated antibodies to assess pathway activation. Numbers under blots represent relative band intensities compared to vehicle control. Representative blots from one of two independent experiments are shown. **I,** Relative *SGK3* mRNA expression, quantified by RT-qPCR, normalized to TBP and to the parental cells in the indicated cell lines. Data represent Mean ± SEM from three independent experiments (*p< 0.05, ***p<0.001, ****p<0.0001, two-tailed t-test). **J,** SGK3 levels in whole cell lysates from the indicated parental cells and lapatinib-DTPs, quantified by immunoblotting. ERK2 serves as a loading control. Numbers under blots represent the normalized relative intensity compared to parental cells. Representative blots from three independent experiments are shown. **K** and **L,** Relative *SGK3* expression, quantified by qRT-PCR, from BT474 (**K**) and HCC1419 (**L**) cells treated with DMSO vehicle (Veh), lapatinib (Lap), fulvestrant (Ful), or lapatinib plus fulvestrant (Lap + Ful) for 48 hours. Expression values were normalized to TBP and to the vehicle control. Mean ± SEM from three independent experiments is displayed. (*p<0.05, **p<0.01, two-tailed t-test).

We next investigated the signaling pathways activated in lapatinib-DTPs that might enable their survival. Despite continuous, complete HER2 inhibition, the PI3K/AKT/mTORC1 and ERK/MAPK pathways were reactivated partially both in luminal-like and in mesenchymal-like lapatinib-DTPs (Supplementary Fig. S7A). However, treatment with pathway-specific inhibitors showed that survival of lapatinib-DTPs required PI3K, but not MEK, activity (Supplementary Fig. S7B-E). Surprisingly, however, treatment with the pan-PI3K inhibitor BKM120 induced higher cytotoxicity than two AKT inhibitors with different inhibitory mechanisms (GSK690693 and MK-2206) tested over a large dose range (Figs. 6E and 6F, Supplementary Figs. S8A and S8B). Compared with BKM120 treatment, GSK690693 and MK-2206 also showed variable and lower, if any, inhibition of phosphorylation of AKT substrates, including PRAS40, TSC2, and GSK3β (Figs. 6G and 6H; Supplementary Figs. S8C and S8D). Furthermore, in all DTPs, mTORC1 activity, as inferred by S6K (T389) and S6 (S240/244) phosphorylation, was inhibited to a far greater extent by BKM120 than by either AKT inhibitor. Moreover, phosphorylation of TSC2 on S939 was inhibited by BKM120, but not by GSK690693 or MK-2206 (Figs. 6G and 6H). These data show that lapatinib-DTPs survive through a PI3K-dependent/AKT-independent mechanism, which nevertheless leads to TSC2 phosphorylation and mTORC1 activation.

### SGK3 is transcriptionally up-regulated by estrogen receptor in luminal-like DTPs

To probe the mechanism of pathway re-wiring in lapatinib-DTPs, we first asked whether activation of this putative pathway depends on PDK1, which phosphorylates the activation loop of multiple AGC kinases (47). Inhibition of PDK1 by GSK2334470 induced cytotoxicity to a similar extent as BKM120 (Supplementary Fig. S8E and S8F), and the cytotoxicity of these inhibitors correlated with their ability to inhibit the phosphorylation of mTORC1 substrates (Supplementary Figs. S8G and S8H). We therefore hypothesized that a PDK1-dependent/AKT-independent kinase was responsible for mTORC1 activation in luminal-like DTPs. *SGK3* is a known estrogen-induced gene in breast cancer, and SGK3 is implicated in AKT-independent survival (48, 49). *SGK3* transcript and its product were up-regulated selectively in luminal-like DTPs (Figs. 6I and 6J), and lapatinib-induced *SGK3* expression in BT474 and HCC1419 cells was blocked by fulvestrant co-administration (Figs. 6K and 6L). Therefore, ER mediates transcriptional up-regulation of *SGK3* in luminal-like DTPs, which in turn activates mTORC1 in a PI3K-dependent/AKT-independent manner.

### SGK3 phosphorylates six sites on TSC2 to activate mTORC1 in an AKT-independent manner

AKT-catalyzed phosphorylation of TSC2 is critical for growth factor-stimulated mTORC1 activation (50–52). Others have shown that SGK1 can phosphorylate TSC2 to activate mTORC1 (53), but whether SGK3 phosphorylates the same sites as AKT and/or which SGK3-evoked phosphorylation sites on TSC2 were responsible for mTORC1 activation was unknown. As expected, recombinant SGK3 phosphorylated TSC2 on AKT substrate motif sites *in vitro* (Fig. 7A). Seven sites on TSC2 (S939, S981, T993, S1130, S1132, T1462, and S1798) conform to the AKT substrate motif, RXRXXS/T. Mass spectrometric analysis of the *in vitro* kinase reaction showed that SGK3 catalyzed increased phosphorylation on six of these (S939, S981, S1130, S1132, T1462, and S1798) compared with the TSC2-only control (Fig. 7B).

**Figure 7.**
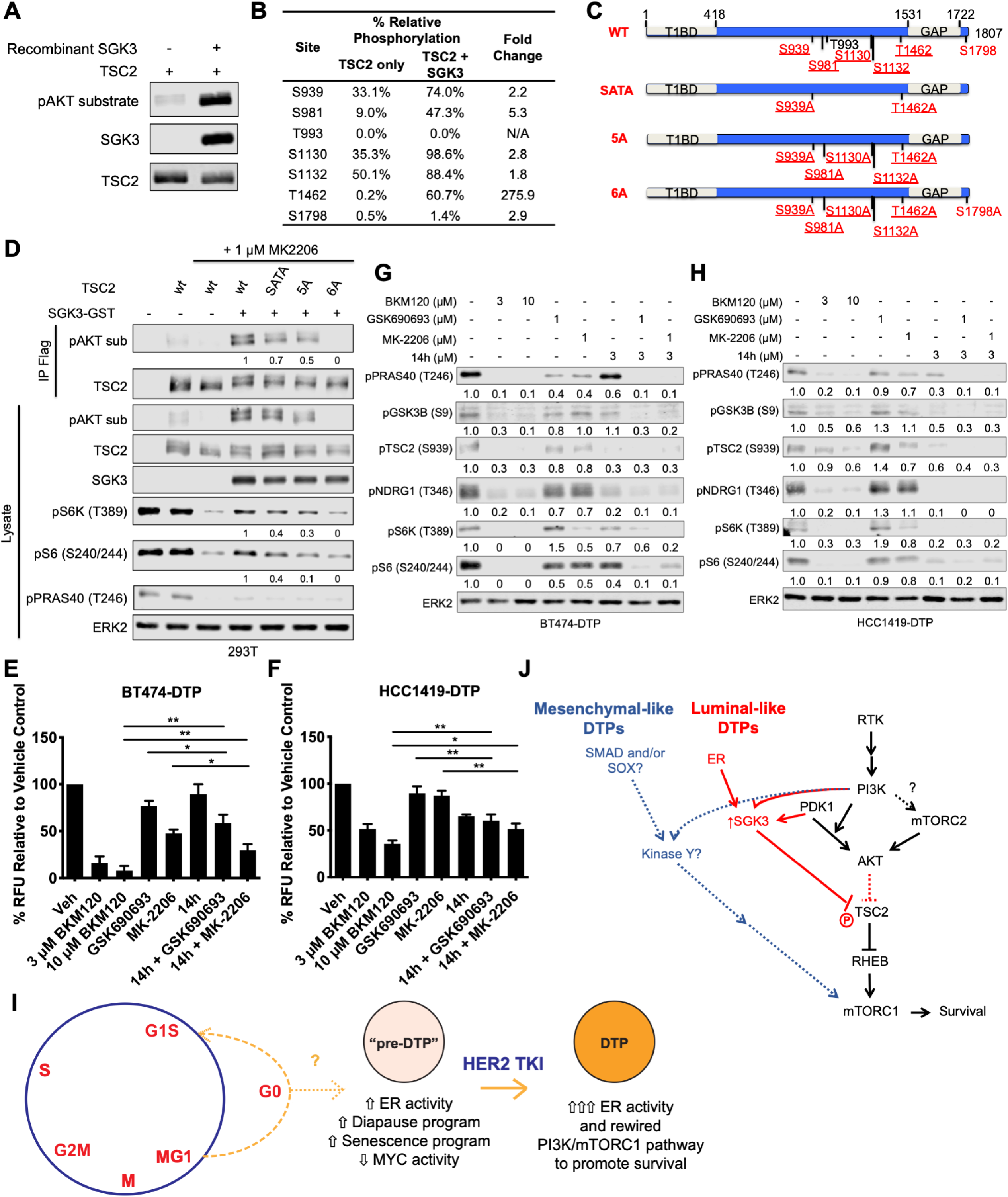
SGK3 phosphorylates TSC2 to activate mTORC1. **A**, FLAG-tagged *TSC2* expression construct was transfected into 293T cells, which were then treated with 1 μM BKM120 for one hour to dephosphorylate PI3K-dependent sites. TSC2 was recovered from cell lysates by immunoprecipitation with ANTI-FLAG M2 agarose beads and incubated with 100 ng recombinant SGK3 at 30°C for 30 minutes in presence of 100 μM ATP. Immunoprecipitates and lysates were subjected to immunoblot analysis with the indicated antibodies. Representative blots from two independent experiments are shown. **B,** A scaled-up *in vitro* kinase reaction was performed as in **A** and analyzed by mass spectrometry. Table shows relative phosphorylation of each pAKT motif site, compared with the TSC2-only control. **C,** Schematic showing wild type (WT) TSC2 and positions of S/T->A TSC2 phosphorylation mutants. Sites conforming to the AKT substrate motif sequence (RXRXXS/T) are displayed with reported AKT phosphorylation sites underlined. SGK3 sites identified by mass spectrometry are highlighted in red. Specific S/T->A mutants analyzed also are shown. **D,** 293T cells were transfected with expression constructs for FLAG-tagged wild type (WT) or mutant *TSC2* with or without *SGK3-GST*, as indicated, and then serum-starved overnight. Where indicated, cells were treated with 1 μM MK-2206 for 30 minutes prior to stimulation with 50 ng/ml IGF1 for 30 minutes, and whole cell lysates and TSC2 immunoprecipitates were immunoblotted with the indicated antibodies. Numbers under the blots represent relative intensities compared with those from cells co-transfected with wt*-TSC2* and *SGK3-GST* + MK-2206; signals from the wt-TSC2 + MK-2206 lanes were subtracted from each before quantification. Representative blots from one of two independent experiments are displayed. **E** and **F**, BT474 lapatinib-DTPs (**E**) and HCC1419 lapatinib-DTPs (**F**) were treated alone or in combination with 3 μM or 10 μM BKM120, 1 μM GSK690693, 1 μM MK-2206, or 3 μM 14h for 96 hours, and cell survival was assessed by Alamar Blue assay. Mean ± SEM of the normalized relative fluorescence units from three independent experiments is displayed (*p<0.05, **p < 0.01, two-tailed t-test). **G** and **H,** BT474 lapatinib-DTPs (**G**) and HCC1419 lapatinib-DTPs (**H**) were treated with the inhibitors for one hour, and whole cell lysates were subjected to immunoblot analysis with the indicated antibodies. Numbers under the blots indicate relative intensity compared with the vehicle control. Representative blots from two independent experiments are displayed. **I,** Proposed model for luminal-like DTP ontogeny. As luminal-like HER2+ breast cancer cells cycle through late mitosis, the population bifurcates, with a subset of cells (M/G1) cycling into G1, while another sub-population transits into G_0_. The latter state (or a subset of cells within this state, indicated by “?”) is “primed” to become DTPs upon TKI exposure. Pre-DTPs also show selective activation of ER target genes and activation/repression of genes associated with DTPs in other systems (diapause, senescence, and MYC targets). Upon exposure to a HER2 TKI, some pre-DTPs further increase their expression of estrogen receptor target genes, including *SGK3*. SGK3 rewires the PI3K/mTORC1 pathway to enable PI3K-dependent, AKT-independent survival. **J,** Distinct survival programs are activated in luminal-like and mesenchymal-like DTPs. In luminal-like DTPs, ER transcriptionally up-regulates *SGK3*, which phosphorylates and inhibits TSC2 to mediate mTORC1 activation and survival. By contrast, mesenchymal-like DTPs exhibit an epithelial-to-mesenchymal (EMT)-like transcriptional program evoked by (an) unidentified transcription factor (s). Mesenchymal-like DTPs also activate mTORC1 in an AKT-independent manner by means of an as yet unidentified kinase.

Mutation of two conserved AKT phosphorylation sites on TSC2 (S939A and T1462A; SATA mutant) prevents most growth factor-stimulated mTORC1 activation (51), although mutation of additional sites (S981, S1130, 1132) is required to completely ablate mTORC1 activity (50,52,54). To ask if these site(s) are required for SGK3-evoked mTORC1 activation, we co-transfected 293T cells with expression constructs for GST-SGK3 alone or with wild type (WT) TSC2 or various phosphorylation site mutants (Fig. 7C). SGK3-evoked mTORC1 activation was assessed in the presence of the AKT inhibitor MK-2206 by immunoblotting with AKT substrate antibodies. The TSC2-SATA and TSC2-5A mutants showed decreased SGK3-evoked phosphorylation, but mutation of all sites phosphorylated by SGK3 *in vitro* was required to abolish SGK3-induced TSC2 phosphorylation in 293T cells (Fig. 7D). Only TSC2-6A completely inhibited SGK3-evoked mTORC1 activation.

As SGK3 is transcriptionally up-regulated in BT474-DTP and HCC1419-DTP, we asked whether SGK3 mediates AKT-independent survival and mTORC1 activation in these cells. We assessed the effects of 14h, a small molecule inhibitor with *in vitro* IC50s of 10 nM and 4 nM against SGK1 and SGK3, respectively (49, 55). Importantly, 14h inhibits SGK1/3 more potently than AKT1 (49). As expected, AKT inhibitors showed partial cytotoxic effects on BT474-DTPs (Fig. 7E). Single agent 14h inhibited survival only slightly, but it showed additive killing when combined with either AKT inhibitor. Nevertheless, 14h/AKT inhibitor combinations still suppressed BT474-DTP survival to a lesser extent than BKM120. AKT or SGK3 inhibition alone only modestly inhibited HCC1419-DTP cell survival (Fig. 7F), but unlike in BT474 cells, combined 14h and AKT inhibitor treatment showed similar killing effects as 14h alone. Notably, SGK3 is up-regulated by 9-fold in HCC1419-DTPs, compared with the HCC1419 parental cells. Conceivably, HCC1419-DTPs are re-wired to become more dependent on SGK3 and thus less responsive to AKT inhibitors.

In parallel, we investigated the effects of 14h and/or AKT inhibitors on AKT substrates and mTORC1 activation. SGK3 inhibition was assessed by monitoring the phosphorylation of its substrate NDRG1 (T346) (56). GSK690693 or MK-2206 showed minimal inhibition, if any, of AKT substrate phosphorylation (Fig. 7G and 7H). Treatment with 14h alone inhibited phosphorylation of PRAS40 (T246), GSK3β (S9), and TSC2 (S939) as well as NDRG1 (T346). Inhibition was more pronounced in HCC1419-DTPs than in BT474-DTPs, consistent with the greater effects of 14h on HCC1419-DTP viability (Fig. 7E and F). In both cell lines, the combination of 14h and AKT inhibitors eliminated phosphorylation of AKT substrates and mTORC1 activation. Knockdown of *SGK3* by siRNA also decreased the PI3K-dependent, AKT-independent phosphorylation of TSC2 and the mTORC1 substrates S6K (T389) and S6 (T240/244), after lapatinib or tucatinib treatment (Supplementary Fig. S9A-S9D). *SGK3* knockdown also reduced the survival of BT474 and HCC1419 cells in response to either TKI (Supplementary Fig. S9E-S9H). In concert, these data show that luminal-like DTPs survive HER2 TKI treatment primarily via ER-induced activation of SGK3, which mediates AKT-independent mTORC1 activation and cell survival.

## DISCUSSION

Most cells in TKI-sensitive cancer cell lines die upon exposure to lethal concentrations of TKIs. By contrast, DTPs survive and can acquire new mutations that confer stable drug resistance and cause disease recurrence (57, 58). Also, during drug holidays, DTPs can revive and acquire stable resistance mutations while proliferating (59). Consequently, understanding DTPs ontogeny could lead to strategies that prevent their emergence and improve disease outcome. We find that upon exposure to the HER2 TKIs lapatinib or tucatinib, HER2+ breast cancer lines give rise to either of two types of DTPs, which have different transcriptomes, survival signaling pathways, and drug vulnerabilities. Barcoding experiments show that HER2 TKI-DTPs emerge stochastically, but fluctuation analysis and scRNA-seq show that, at any given time, a fraction of untreated cells exist in a “pre-DTP” state, characterized by a G_0_-like signature and expression of a subset of DTP genes, and are primed to become DTPs (Fig. 7I). Pre-DTPs purified from bulk parental cells by FACS for NPY1R or ABCC5 surface expression exhibit enhanced ability to become DTPs upon HER2 TKI exposure, providing direct functional evidence for the pre-DTP state. Luminal-like DTPs activate a subset of ER target genes, but upon progression to DTPs, ER activity is induced further and *SGK3* transcription is upregulated. SGK3 then phosphorylates and inactivates TSC2 to mediate mTORC1 activation and survival (Fig. 7J). Mesenchymal-like DTPs do not induce ER activation, yet they also activate mTORC1 in an AKT-independent fashion, via an undefined kinase(s) (Fig. 7J). Most importantly, cells resembling pre-DTPs are seen in HER2+ breast tumors and appear to correlate with decreased response to neo-adjuvant lapatinib treatment. Our results provide new insights into HER2 TKI-DTP ontogeny and identify potential vulnerabilities for these cells. We also provide evidence that “pre-DTPs” might also exist in other sub-types of breast cancer.

Previous studies showed that some HER2+ breast cancer lines could give rise to lapatinib-DTPs, but whether this was a general property of such lines remained unclear (21,32,60). Nearly half of the HER2+ breast cancer lines that we examined (Type III lines) displayed characteristic DTP behavior. By contrast, others (Type I/Type II lines) proliferated or failed to die upon TKI exposure. All Type I/II cells have *PTEN* copy number loss or “hotspot” activating *PIK3CA* mutations (i.e., E545K or H1047R), and PI3K pathway activation can cause resistance to lapatinib (13,61,62). However, two Type III lines (BT474, EFM192A) also harbor potentially transforming *PIK3CA* mutants (K111N, C420R) (39), so the mere presence of such a mutation does not preclude DTP generation. Notably, Vasan *et al.* demonstrated that mutant *PIK3CA* alleles differ in transforming potency in human breast epithelia, although they did not study these particular mutants (63).

Contrary to Sharma *et al*.’s initial report on NSCLC (21) and other studies (32,33,58,64), only one Type III line (EFM192A) gave rise to DTEPs. The NSCLC lines studied by Sharma *et al.* and others contain cells with pre-existing TKI resistance mutations, which give rise to “early resistance”. By contrast, 12-16 weeks of TKI exposure are needed before “late resistance” emerges (57). Conceivably, our 8-week observation period might have been too short to observe the latter. Two very recent studies identified cells with properties of “cycling DTPs” in several cancer cell lines (32, 33); presumably, these cells correspond to DTEPs. However, consistent with our failure to observe cells with DTEP-like behavior in most Type III lines, cycling DTPs were not observed in the two HER2+ breast cancer lines analyzed by Oren *et al*. (32).

HER2+ tumors can be divided into “HER2-enriched” and “HER2-luminal” subtypes (65). The former express higher levels of RTK and mesenchymal genes; the latter feature luminal and estrogen response genes (66). We find that cell lines that model these patient subgroups yield DTPs with distinct transcriptomes. Luminal-like DTPs survive via an ER-driven program, as demonstrated by their sensitivity to HER2-TKI/fulvestrant combinations. These results provide a mechanistic underpinning for the empirically derived use of anti-estrogen plus HER2-targeted therapy in HER2+/ER+ patients (67). By contrast, mesenchymal DTPs appear to use a SMAD/SOX-driven transcriptional program. It will be important to delineate the molecular drivers for this program (e.g., TGFβ, BMPs) so that it too can be targeted prospectively in patients with HER2+/ER-tumors. While our manuscript was in revision, diapause (29, 30), senescence-like (31), chemotherapy-induced (31), and MYC-gene signatures (29–31), were found to be upregulated or repressed in other DTPs. We also had noted downregulation of MYC genes in our original analyses, but the other signatures were not present in the pathway gene sets that we initially used for GSEA. Re-analysis with an expanded collection revealed enrichment for the diapause and, in some cell lines, the chemotherapy-induced and senescence-like signatures. Thus, while luminal and mesenchymal programs dominate the transcriptomes of DTPs from HER2+ER+ and HER2+ER-lines, respectively, these cells also share features with DTPs induced by other agents and in other tumor types. Notably, these signatures are also detectable in select cells from primary HER2+ and triple negative breast cancers.

Concepts such as dormancy, quiescence, “cancer stem cells”, persistence, residual disease and their relationship to disease recurrence and therapeutic resistance have long been debated (68–71). The classical “cancer stem cell” (CSC) hypothesis, for example, posits a defined hierarchy in which a limited number of CSCs can self-renew and give rise to all other cells in a tumor (72, 73). CSCs are often portrayed as slow-cycling and more drug-resistant than bulk tumor cells (although these properties are not necessarily intrinsic to the concept). Earlier studies of other malignancies, using limiting dilution, cell surface marker, and/or genetically encoded reporter approaches (22,24,74), suggested that, by contrast, DTPs have a stochastic origin. Our lentiviral barcoding experiments provide unambiguous evidence that HER2 TKI-DTPs arise stochastically: over a several week period, essentially every initially tagged HER2+ breast cancer cell exhibited the capacity to give rise to a DTP. While our manuscript was in revision, two groups used lentiviral barcoding of xenografts to reach similar conclusions about chemotherapy-induced DTPs (29, 30). In concert, then, the preponderance of the evidence argues against drug resistance arising from intrinsically therapy-resistant CSCs in HER2+ breast cancer and most, if not all, other malignancies.

Yet while every cell can, *over time*, become a DTP, fluctuation analysis implies that at *any given time*, HER2+ breast cancer cells differ in their propensity to become DTPs. Consistent with this implication, scRNA-seq reveals “pre-DTPs”, characterized by a G_0_-like transcriptional signature, increased or decreased expression of subsets of DTP DEGs that are up- or down-regulated, respectively, in DTPs, and consistent repression of MYC genes and genes that are downregulated during embryonic diapause. Trajectory analysis of two HER2+/ER+ cell lines indicates that G_0_-like pre-DTPs give rise to most DTPs, and direct evidence is provided by the increased DTP-forming activity of untreated BT474 cells expressing high levels of either of two cell surface markers suggested by the scRNA-seq analysis to be enriched in pre-DTPs, NPY1R or ABCC5. NPY1R^hi^-enriched BT474 cells have increased basal expression of ER target genes and further induce the ER transcriptome upon TKI exposure. These findings are consistent with a model in which a subset of cells (pre-DTPs) is “primed” for induction into *bona fide* DTPs upon TKI (and possibly, chemotherapy) exposure, as opposed to the concept of drug-induced epigenetic change. This “priming/induction” model implies that some, although certainly not all, genes enriched in DTPs reflect selection for pre-DTP genes. For example, Hangauer *et al.* reported that compared with parental cells, BT474-DTPs feature global downregulation of antioxidant genes, including genes encoding glutathione peroxidases (e.g., *GPX1, GPX2, GPX4*). They also found that GPX4 inhibition is selectively toxic to DTPs. BT474 pre-DTPs also display lower levels of *GPX1* and *GPX4* compared with other untreated cells (Supplementary Tables 4 and 5). These findings suggest selection of pre-existing GPX1/4-low pre-DTPs, rather than induction of a GPX1/4-low state and provide an alternate explanation for why GPX4 inhibitor pre-treatment prevents lapatinib-DTP generation (66).

Our results indicate that nearly all (∼90%) BT474-DTPs arise from cells with a G_0_-like expression signature (DTP1). Although it is generally believed that cultured cells in complete media cycle continuously and enter G_0_ only upon growth factor depletion, Spencer and Meyer showed that depending on the level of CDK2 activity at M phase exit, cells transit either through G_0_ or directly enter G_1_ (75). Our finding that untreated HER2+ breast cancer lines harbor cells with a G_0_-like transcriptome extends their observations. *BRAF^V600E^*-mutant melanoma cell lines (especially in 3D culture) also have a “slow cycling” fraction of cells marked by JARID1B (*KDM5B*) expression, which is required for continuous passaging of tumors (74). Similar to the pre-DTPs in our study, JARID1B-positive and -negative cells stochastically interconvert, and vemurafenib treatment enriches for JARID1B+ cells (25). These lines also stochastically express select RTKs (e.g., EGFR, TRKA/NGFR, AXL) in the absence of drug exposure, and these RTK+ cells are enriched for vemurafenib resistance (24). Human melanoma samples also have small populations of *KDM5B*-positive cells (74) and scRNA-seq (76) shows that *KDM5B* expression correlates with a “G_0_/G_1_” signature. Furthermore, cells in a G_0_-like neural crest state pre-exist in a zebrafish melanoma model and are selected for upon BRAF inhibitor treatment (77). While none of these reports explicitly equate G_0_-like cells with pre-DTPs, in concert these data are certainly consistent with such a model.

Conversely, a small fraction of DTPs (DTP2) is enriched for G2/M genes (Fig. 5K). Conceivably, DTP2 cells correspond to the recently observed “cycling DTPs” (32, 33). Because we saw no expansion/colony formation of lapatinib- or tucatinib-treated BT474 cells over an ∼8-week period, we cannot exclude the possibility that these cells are growth arrested permanently or committed to die. Interestingly, the DTP1 and DTP2 states correspond to the two “decision” points in the cell cycle (G_0_/G_1_ and G_2_/M) identified by Spencer and Meyer, raising the possibility that cellular plasticity might be greatest in these cell cycle phases. Interestingly, *Drosophila* neural stem cells subjected to nutritional deprivation (the same conditions that induce diapause) arrest in, and re-enter the cell cycle from, either G_0_ or G_2_/M (78). Regardless, “non-cycling” DTPs comprise the majority DTP population in HER2+ breast cancer lines (as well as in nearly all cell lines studied in these recent reports), remain capable of resuming proliferation, and forming new tumors after drug removal, and appear to be present in primary HER2+ tumors from patients. Furthermore, a high G_0_ expression signature correlates with decreased pCR in a clinical trial of neoadjuvant lapatinib. Hence, strategies to eliminate both non-cycling and cycling DTPs will be required to obtain durable remissions/cures.

Our results suggest several such strategies. As determined empirically in the clinic, combining fulvestrant with HER2 TKI results in markedly decreased DTP formation. Notably, BCL2 is up-regulated in luminal-like DTPs (Supplementary Table 2) and might be targeted with BH3 agonists (e.g., Venetoclax). Furthermore, we find that luminal- and mesenchymal-like DTPs reactivate PI3K/mTOR signaling via pathway rewiring. Although combining lapatinib and PI3K inhibitors could be limited by toxicity (79), targeting these rewired pathways could prove beneficial. Comporting with previous reports that SGK1 or SGK3 can mediate AKT-independent activation of mTORC1 and survival in cancer cells treated with PI3K⍺-specific or AKT inhibitors (49, 80), we find that *SGK3* is a key ER target that mediates survival in luminal-like DTP and could be targeted to prevent DTP generation. Our results also provide new mechanistic insight into how SGK3 activates mTORC1. AKT phosphorylates five sites on TSC2 (S939, S981, S1130, S1132, T1462), and all of these sites are required for AKT-mediated mTORC1 activation (50–52,54). Our MS analysis demonstrates that SGK3, like SGK1 (80), can phosphorylate these five sites on TSC2, as well S1798. Furthermore, all six SGK3-evoked sites must be ablated to abolish SGK3-mediated mTORC1 activation. Our results also contrast with previous work arguing that PRAS40 is a selective AKT target (49); at least when SGK3 levels are increased, it can mediate PRAS40 phosphorylation (Fig. 7H). Conversely, NDRG1 (T346) phosphorylation is primarily dependent on SGK3, although we did observe additive inhibition of NDRG1 (T346) phosphorylation when AKT and SGK3 inhibitors were combined, consistent with previous reports that both kinases can phosphorylate this site (80–82).

While mesenchymal DTPs also rewire their signaling pathways to enable AKT-independent mTORC1 activation, the alternative kinase and detailed mechanism remains to be elucidated. Future work is also needed to uncover the epigenetic mechanism(s) for priming of the “pre-DTP” state and its transient, but differential heritability in HER2+ breast cancer cells.

## MATERIALS AND METHODS

### Reagents

Tissue culture reagents, including regular DMEM, RPMI, and FBS were purchased from Wisent Bioproducts. PD0325901 was synthesized as described (83), and 14h was synthesized by BioDuro according to a published protocol (55). Lapatinib ditosylate was purchased from LC Laboratories, tucatinib was purchased from MedChemExpress, 5-FU was purchased from Sigma, and Trastuzumab was obtained from the NYU Langone Health pharmacy. Fulvestrant, BKM120, MK2206, and GSK690693 were purchased from Selleck Chemicals. GSK2334470 was purchased from Tocris. Recombinant SGK3 (cat #14-647) was purchased from EMD-Millipore.

### Plasmids and Site-directed Mutagenesis

The expression construct for *SGK3-GST* was provided by Dr. Alex Toker (Beth Israel Deaconess Medical Center, Boston, MA). pcDNA3-based plasmids encoding FLAG-tagged wild type and SATA (S939A/T1462A)-mutant *TSC2*(51) were obtained from Addgene. The *TSC2-5A* (S939A, S981A, S1130A, S1132A, T1462A) and *TSC2-6A* plasmids (S939A, S981A, S1130A, S1132A, T1462A, S1798A) were constructed using *TSC2*-SATA as the template for site-directed mutagenesis and the QuikChange Multi Site-Directed Mutagenesis Kit (Agilent Technologies). The sequences of all point mutations were verified by Sanger sequencing.

### Cell Culture and Transfections

Cell lines were purchased from the American Type Culture Collection or Deutsche Sammlung von Mikroorganismen und Zellkulturen, and their genotypes were confirmed by short tandem repeat (STR) analysis (84). Cells were tested regularly for mycoplasma by using a PCR-based kit from Agilent Technologies. MDA-MB-453, BT474, SKBR3, UACC893, and 293T cells were maintained in DMEM supplemented with 10% fetal bovine serum (FBS) and penicillin and streptomycin (Pen/Strep). MDA-MB-361 cells were maintained in DMEM supplemented with 20% FBS and Pen/Strep. HCC1569, HCC202, HCC1419, and EFM192A cells were maintained in RPMI supplemented with 10% FBS and Pen/Strep. SUM225 cells were maintained in Ham’s F12 supplemented with 5% FBS, 5 μg/ml insulin, 1 μg/ml hydrocortisone, and 10 mM HEPES. Transient transfections of plasmids were performed by using LipoD293™ In Vitro DNA Transfection Reagent (SignaGen Laboratories). Transient transfections of SMARTPool SGK3 siRNA (L-004162-00-0005, Horizon Discovery) were performed with Lipofectamine RNAiMAX Transfection Reagent (Thermo Fisher Scientific).

### Cell Proliferation Assays

HER2+ breast cancer lines were exposed to 2.5 μM lapatinib or 1.2 μM tucatinib. Viable cell number at each time point was quantified by using a Vi-Cell counter (Beckman-Coulter), and % survival was calculated relative to viable count before drug exposure (t=0). To assess the reversibility of lapatinib/tucatinib tolerance, cells were trypsinized, washed three times in PBS, and re-plated in standard growth media without lapatinib/tucatinib. At the indicated times after TKI withdrawal, cells were trypsinized and re-plated on 6-well plates at 500,000-1,000,000 cells/well. Cells were allowed to attach overnight, some wells were trypsinized to obtain an initial viable cell count (Count_t0_), and the rest were treated with lapatinib for 7 days to obtain the day 7 viable cell count (Count_t7_). Survival of cells following drug re-challenge was calculated as Count_t7_/Count_t0_.

For assessing the response of parental cells to lapatinib in combination with fulvestrant or estrogen depletion, cells (10,000/well) were seeded into 96-well plates. NPY1R-selected cells were treated immediately after FACS with lapatinib and/or the indicated treatments for 14 days and then switched to the regular growth media without the drug for 14 days to assay for regrowth. Residual cells were imaged by using an IncuCyte apparatus with a colored mask used to show cells that remained on the plate. ABCC5-selected cells were treated immediately after FACS with lapatinib or tucatinib for 14 days. Cells were then counted using Countess II FL Automated Cell Counter (Invitrogen) with trypan blue to exclude dead cells.

The cytotoxic effects of various drugs were assessed by treating DTPs in 96-well plates (1,000-3,000 cells/well) with serial dilutions of each agent for 96 hours. At the assay end point, cell number was estimated by using AlamarBlue® Cell Viability Reagent (Life Technologies) and measuring fluorescence (excitation: 540 nm; emission: 590 nm) with a Spectramax microplate reader (Molecular Devices).

### Tumorigenicity Assays

The indicated numbers of parental BT474 cells and BT474-DTPs in 1:1 HBSS:Matrigel (BD Biosciences) were injected into the mammary fat pads of NSG mice. Mice were monitored for tumor formation for up to 5 months.

### Lentiviral Barcoding

#### Barcode library

Oligonucleotides comprising a 12 base pair degenerate region (the barcode) followed by two stable bases (C or G) and one of several four base pair library codes were synthesized with common flanking regions (Sigma Aldrich, St. Louis, MO, USA). Nested PCR using the common regions generated double-stranded DNA, which was ligated into the second-generation lentiviral vector pLJM1, which contains a puromycin resistance cassette and ZsGreen fluorescent marker. Three barcode libraries, each identifiable by a unique library code, were cloned, transformed into *E. coli*, and plated as a pool on solid media. More than 5×10^6^ bacterial colonies were scraped and pooled for two of the high diversity libraries. Plasmid DNA was isolated, and a sample was sequenced to confirm a diversity of >10^6^ unique barcodes. These libraries were named Library 0 and Library 1. The third library was generated to use for standard spike-in controls. Single colonies were selected, prepped and Sanger sequenced to identify several standard barcodes.

#### Lentiviral Transduction and Barcoding Experiments

Lentiviruses containing barcode plasmid libraries were produced in HEK293T cells. BT474 cells (1×10^6^) were plated on a 6 cm dish and infected with the lentiviral library at MOI=0.1 to ensure that each barcode was present in only a single cell. The infected population was expanded to 50 million cells, and cells were not discarded during passaging to preserve representation of the barcodes. One million cells were analyzed in triplicate to assess initial representation of the barcodes. Ten million cells were plated in triplicate on 15 cm dishes, and cells were treated with 2.5 µM lapatinib for 14 days (experiment 1). The remaining cells were kept in culture until the start of experiment 2 (below). After 14 days of lapatinib treatment, the remaining cells (approximately 1 million cells) were collected for genomic DNA (gDNA) extraction. The experiment was repeated (experiment 2) after 14 days of culture of the remaining cells from above. Single barcoding experiments were performed on HCC1419, SKBR3, and EFM192A cells by following the protocol described above for experiment 1.

#### Barcode amplification and sequencing

The gDNA for all samples was adjusted to 400ng/µL using nuclease-free water. Sequencing libraries were constructed by PCR amplification using a common 3’ primer “BL Seq Amp 3’: AATGATACGGCGACCACCGAGATCT and one of 166 unique 5’ primers “BL Seq Amp 5’ XXX”:CAAGCAGAAGACGGCATACGAGATNNNNNNCGATTAGTGAACGGATCTCGA CGGT, where the “N”s represent a unique sample index. Each gDNA was amplified as a technical triplicate with unique indexes using ExTaq (Takara Cat#RR001A) with PCR program of 95°C for 5 minutes, 94°C for 30 seconds, 65°C for 30 seconds, 72°C for 30 seconds, and back to step 2, 32x followed by a 5 minute hold at 72°C. PCR efficiency was assessed by running the product on a 3% agarose gel. The 137 bp barcode library band was quantified using Bio Rad Image Lab software. Equal amounts of each PCR product were pooled into batches and purified on 15% TBE PAGE gels (Novex). Purified PCR products were quantified by using a Qubit, pooled, and sequenced on an Illumina HiSeq2500 with version 4 chemistry using Illumina sequencing primer: ACACTCTTTCCCTACACGACGCTCTTCCGATCT and custom primer: ATCGATACCGTCGAGATCCGTTCACTAATCG for multiplexed sample ID. Samples were de-multiplexed and barcode abundances were analyzed.

#### Barcode Processing and Analysis

FASTQ files for each sequenced sample were processed using a bespoke Perl script. Each read was examined to identify one of the three expected library codes (CCAA, ACGT, or TGGA) followed by eight bases corresponding to the vector sequence (e.g., ATCGATAC), allowing up to one mismatched base for each feature. Reads lacking both of these sequences were discarded. The nucleotide sequence corresponding to the barcode was extracted as the 18 nucleotides preceding the vector sequence, and all unique barcodes were counted. All barcode count files, one per sample, were then merged into a single matrix. Noise introduced through sequencing or PCR errors was reduced by collapsing barcodes within a Hamming distance of two into a single barcode record, where the barcode with the highest average abundance was retained as the “parent” barcode. Next, samples with fewer than 100,000 filtered sequence reads were removed, and each sample was normalized for sequencing depth by dividing all read counts in a sample by the sum of all read counts. Technical replicates were combined by averaging, and barcodes that were observed in only one sample were removed as potential artefacts. The final barcode matrix comprised 154,262 barcode sequences.

#### Fluctuation analysis

We consider a model wherein individual cells switch stochastically between the non-DTP and a DTP state. Each state is transiently heritable; i.e., cells remain in that state for multiple generations before switching to the other state. Recent work used the classical Luria-Delbrück fluctuation test framework to estimate switching rates based on variations in the number of DTP-like cells between single-cell derived lineages (24, 43). In essence, if DTP cells arise purely randomly, then one expects a Poisson distribution (i.e., minimal fluctuation) for the number of DTP cells across lineages. By contrast, large lineage-to-lineage fluctuations imply transient heritability of the DTP state with increasing heritability driving enhanced fluctuation.

Let f denote the fraction of cells that are DTPs. Then, the extent of fluctuation in the fraction of DTP cells between lineages (as quantified by the coefficient of variation CV) is given by:

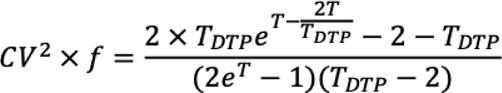

where T is the number of generations that single cells were expanded before treatment, and T_DTP_ is the average number of generations a cell remains in the DTP state before switching to a non-DTP state (43). Based on the barcoding data, we first compute the fraction of DTP cells in each lineage by taking the ratio of the number of reads after 14-days of treatment to that at the start of treatment. The analysis was restricted to lineages with >5,000 barcode counts at treatment onset. We obtain CV^2^ = 0.96, 0.13, 2.2, 0.25 for EFM192A, HCC1419, SKBR3 and BT474, respectively. To estimate the technical noise, we quantified variations in the fraction of DTP cells between technical replicates averaged across all lineages and found this variation to be ∼0.04. As the technical noise is at least an order of magnitude lower than the computed inter-lineage CVs, this result strongly argues against the random Poisson model and points to memory in the DTP state. We then use the above equation to calculate transient heritability (T_DTP_) from the CV values. For these experiments, t = 6 generations, and f = 0.1 for SKBR3 and BT474, f = 0.3 for EFM192A, f = 0.4 for HCC1419 (see Fig. 1A). Hence, the transient heritability of the DTP state is estimated as 6.2 ± 0.9 (SKBr3), 2.6 ± 0.4 (HCC1419), 7.3 ± 1 (EFM192A), and 2.2 + 0.4 (BT474), where the 95% confidence interval is estimated using bootstrapping. The experimentally obtained distributions of the fraction of DTP cells together with model fits are illustrated in Supplementary Fig. S2C. Model fits are based on a beta distribution with mean f and coefficient of variation as predicted by the above equation.

### Bulk RNA-seq

Bulk RNA sequencing was conducted by the Princess Margaret Genomics Center or the Perlmutter Cancer Center (PCC) Genome Technology Center (GTC). RNA was extracted using the RNeasy Mini Kit followed by treatment with RNase-free DNase to remove contaminating genomic DNA. RNA (200 ng) was reverse transcribed by using the Illumina TruSeq Stranded Total mRNA kit. Libraries were sized on an Agilent Bioanalyzer, and their concentrations were validated by qPCR. The libraries were then loaded onto an Illumina NextSeq cartridge V2 for cluster generation, and the flow cell was subjected paired-end sequencing on Illumina Nextseq500. For lapatinib DTPs, alignment was performed with STAR v.2.5.2, and reads were mapped to the human reference genome (hg38). Mapped reads were quantified by RSEM v.1.3.0, and differential expression was assessed by using DESeq2. DEGs were identified by a 2×2 analysis that compared genes expressed in DTP and parental cells after separating lines into luminal-like (BT474 and HCC1419) and mesenchymal-like (EFM192A and SKBR3) groups. Sequencing reads for tucatinib DTPs were mapped to the reference genome (hg19) using the STAR aligner (v2.5.0c) (85). Alignments were guided by a Gene Transfer Format (GTF) file. Mean read insert sizes and their standard deviations were calculated using Picard tools (v.1.126) (http://broadinstitute.github.io/picard). The read count tables were generated using HTSeq (v0.6.0) (86), normalized based on their library size factors using DEseq2 (87), and differential expression analysis was performed. The top 500 differentially expressed (sorted by the highest adjusted p value) genes were visualized in heatmaps. The Read Per Million (RPM) normalized BigWig files were generated using BEDTools (v2.17.0) (88) and bedGraphToBigWig tool (v4). Gene set enrichment analysis was performed using GSEA tool (89). Samples were compared by principal component analysis or Euclidean distance-based sample clustering. All downstream analyses were performed in R environment (v3.1.1) (https://www.r-project.org/). ChIP Enrichment Analysis (ChEA) was performed with Enrichr. Pathway and gene ontology analysis were performed by ranking genes according to fold-change in the 2×2 or the 4×4 analysis using Bader lab pathway gene sets (Human_GOBP_AllPathways_no_GO_iea_August_01_2018_ symbol.gmt, http://download.baderlab.org/EM_Genesets/August_01_2018/Human/symbol/). To compare the overlap of DEGs between DTPs and NPY1R^hi^ cells, the cut-off was set at abs FC>2 and p<0.05. Statistical enrichment was then assessed by calculating the Fisher’s exact test. To ask if NPY1R^hi^ breast cancers are enriched for ER targets, transcript enrichment was assessed by using cBioPortal and TCGA human breast cancer datasets (65).

### Analysis of NeoALTTO trial data

RNA-seq data on tumor samples from the NeoALTTO clinical trial (90) were processed using STAR aligner (85) and the R package Rsamtools (version 1.24.0). G_0_ gene signatures along with the gene specific weight of +1 or -1, indicating the direction of association with the G_0_ state, were obtained from the original publication using FDR q<0.1 (91). These gene-specific weights along with z-score normalized count per million values were used to compute the weighted sum score for each sample in the dataset. Association between weighted gene signature score and pathologic complete response (pCR) was computed using two-sided Student’s t-test. An analogous approach was used to compute the diapause, CISG, senescence, and Myc targets scores in Supplementary Fig. S4.

### Single Cell RNA-sequencing

BT474 and HCC1419 samples were prepared by the Princess Margaret Genomic Centre following the 10X Genomics Single Cell 3’ Reagent Kits v2 user guide. Briefly, samples were washed two times in PBS (Life Technologies) + 0.04% BSA (Sigma), re-suspended in PBS + 0.04% BSA, and loaded onto a 10X single cell A chip. After droplet generation, samples were transferred onto pre-chilled 96-well plates (Eppendorf), heat-sealed, and incubated overnight in a Veriti 96-well thermal cycler (Thermo Fisher) for reverse transcription (RT). Following RT, cDNA was recovered using the Recovery Agent provided by 10X and purified by using Silane DynaBead (Thermo Fisher) mix, as outlined by the user guide. Purified cDNA was amplified for 13 cycles before purification on SPRIselect beads (Beckman). Samples were diluted 4:1 (elution buffer (Qiagen):cDNA), and cDNA concentration was determined with a Bioanalyzer (Agilent Technologies). cDNA libraries were prepared as outlined by the Single Cell 3’ Reagent Kits v2 user guide with modifications to the PCR cycles based on the calculated cDNA concentration.

The molarity of each library was calculated based on library size, as measured by a Bioanalyzer and qPCR quantification (Roche/Kapa BioSystems). Samples were pooled and normalized to 10 nM, then diluted to 2 nM using elution buffer (Qiagen) containing 0.1% Tween20 (Sigma). Each 2 nM pool was denatured in an equal volume of 0.1N NaOH for 5 minutes at room temperature. Library pools were further diluted to 20 pM using HT-1 (Illumina), before dilution to a final loading concentration of 16 pM, and 100 μl from the 16 pM pool were loaded into each well of an 8-well strip tube and placed onto a cBot (Illumina) for cluster generation. Samples were sequenced on a HiSeq 2500 V4 with the following run parameters: Read 1 – 26 cycles, read 2 – 98 cycles, index 1 – 8 cycles.

EFM192A, SKBR3, SUM225, and primary breast tumor samples were sequenced by the GTC. Primary samples were processed as previously described (92). Cell lines were stained using Biolegend TotalSeq anti-human Hashtag Antibodies 7-10 (Cat #’s 394613, 394615, 394617, 394619) following the New York Genome Center Technology Innovation Lab’s posted protocol (https://citeseq.files.wordpress.com/2019/02/cell_hashing_protocol_190213.pdf). Stained cellular suspensions were loaded on a 10x Genomics Chromium instrument to generate single-cell gel beads in emulsion (GEM). Approximately 10,000 cells/hash-tagged population were loaded per channel. Single-cell RNA-Seq and hashtag libraries were prepared using the following Single Cell 3’ Reagent Kits v3.1: Chromium Next GEM Single Cell 3’ GEM, Library & Gel Bead Kit v3.1, PN-1000121; Chromium Next GEM Chip G Single Cell Kit, PN-1000120; Chromium Single Cell 3’ Feature Barcode Library Kit, PN-1000079 and Single Index Kit T Set A PN-1000213 (10x Genomics), as described in (93) and the Single Cell 3’ Reagent Kits v3.1 User Guide with Feature Barcoding Technology (Manual Part # CG000206 Rev D). Libraries were sequenced on an Illumina Nextseq 2000 using paired-end reads, read1 was 28 cycles, i7 index was 8 cycles, and read2 was 91 cycles. The libraries comprised 50% of the flow cell yielding >49% sequencing saturation.

### scRNA-seq Data Analysis

#### Pre-processing

After confirming cDNA integrity, library quality, number of cells sequenced, and mean number of reads per cell, Cell Ranger Single-Cell Software (https://support.10xgenomics.com/single-cell-gene-expression/software/pipelines/latest/what-is-cell-ranger) was used to map the reads and generate gene-cell matrices. Quality control was performed to calculate the number of genes, UMIs, and the proportion of mitochondrial genes for each cell using the iCellR R package (v1.6.4) (https://cran.r-project.org/web/packages/iCellR/index.html). Cells with particularly low or high numbers of covered genes were filtered. The matrix was normalized (LogNormalize) by dividing the feature counts for each cell by the total counts for that cell and multiplying by the scaling factor and then the matrix was log transformed (log1p). Highly expressed and dispersed genes were used as a gene model for PCA. To fine-tune the results, a second round of PCA was performed based on the top 20 and bottom 20 genes predicted in the first 10 dimensions of PCA (400 genes). Uniform Manifold Approximation and Projection (UMAP) and clustering were performed on the top 10 PCs.

#### Cell Cycle Analysis

The cell cycle phase-specific gene signatures defined by Xue *et al.* (45) were used to calculate phase-specific scores for G_0_, G1S, G2M, M, MG1, and S. To account for differences in gene set sizes, we used a scoring method similar to Tirosh *et al.* (76), implemented in the “i.score” function of iCellR. For Figure 3, we compared the sum of phase-specific gene expression (log10 transformed UMIs) to the distribution of random background gene sets, where the number of background genes is identical to the phase-specific gene set and are drawn from the same expression bins. Each cell was assigned to a cell-cycle stage based on its highest phase-specific score. Cell-cycle UMAP was performed using all of the cell-cycle-phase-specific gene-expression signatures as the input features in the RunUMAP function (umap.method = “umap-learn”, metric = “correlation”). For Figure 4, the raw data was normalized as described above using “LogNormalize”, and the scores were calculated with the following settings: i.score(my.obj, scoring.List = c(“G0”,“G1S”,“G2M”,“M”,“MG1”,“S”), scoring.method = “tirosh”, return.stats = TRUE). Each cell’s cell cycle phase was assigned as its highest cell cycle score. Scoring for other signatures was performed by using the same approach, and scores were then displayed as boxplots. DTP gene expression signature scores were calculated by selecting the top 150 most significant differentially expressed genes from bulk RNAseq and applying the approach for calculating the cell-cycle-phase specific score described above. All gene signatures are in Supplementary Table S12.

#### Trajectory analysis

After pre-processing of the scRNA-seq data, we selected the first 15 dimensions of the PCA for each time point (untreated, 6h-treated, DTP) of BT474 and HCC1419 cells for additional clustering and dimensionality reduction. Clustering was performed by using iCellR with the following settings: iCellR options: clust.method = “ward.D”, dist.method = “euclidean”, index.method = “kl”; phonograph options: k =200, dims = 1:15 on these principal components, and dimension reduction was accomplished by using Uniform Manifold Approximation and Projection (UMAP) (94). For each cell line, we used Monocle2 (v2.16.0) to define pseudotime trajectories. The raw count matrix was used as the input, and the first 10 PCA dimensions of genes identified by iCellR were used for building the trajectories. The reduceDimension function with “DDRTree” algorithm was used to build the trajectory. Cells in the same branch of the trajectory share similar gene expression patterns.

#### Visualization and Statistical Analysis

All visualization and statistical analyses were performed in R (v4.0.0) (http://www.r-project.org/). Two groups were compared using an unpaired t-test. Significance was defined as ns, P > 0.05; *, P < 0.05; **, P < 0.01; ***, P < 0.0001; ****, P < 0.0001. Heat maps were generated by using the pheatmap package (v1.0.12) with scale = “row” argument.

### Quantitative RT-PCR

RNA was extracted using the RNeasy Mini Kit (Qiagen), mRNAs were reverse transcribed with SuperScript III First Strand Synthesis (Invitrogen), and iQ™ SYBR® Green Supermix was used for qPCR on CFX96 (Bio-Rad). Relative expression was calculated with the delta-delta Ct method (ΔΔCt). Briefly, the Ct value of the gene of interest was subtracted from the Ct value of the housekeeping gene TBP to yield ΔCt and the ΔCt from the sample of interest is subtracted with control sample to yield ΔΔCt. The relative expression is expressed as 2^-ΔΔCt^. The following primers were used for qPCR: SGK3: F, 5’-GTGCCCGAAGGTTGCATGAT-3’ and R, 5’-ATCCCTCAAGAGCACACCAA-3’; NPY1R: F, 5’-GCAGGAGAAATACCAGCGGA-3’ and R, 5’-TCCCTTGAACTGAACAATCCTCTT-3’; TBP: F, 5’-TGTGCACAGGAGCCAAGAGT-3’ and R, 5’-ATTTTCTTGCTGCCAGTCTGG-3’.

### Flow Cytometry and Fluorescence Activated Cell Sorting (FACS)

Cells were released from plates by incubating with 0.05% trypsin and then stained with primary antibody (1 μg/10^6^ live cells) against NPY1R (MAB6400; R&D Systems) followed by secondary antibody against human IgG for 30 minutes on ice and/or with PE-conjugated ABCC5 (sc-376965 PE; Santa Cruz). Flow cytometry was conducted on a Becton Dickinson analyzer, and FACS was performed by using a FACSAriaII. Post-FACS purity checks were performed to ensure that each fraction was >90% of the population of interest. Where specified, RT-qPCR was conducted to determine whether the sorted fractions showed the expected differential expression of *NPY1R*. For NPY1R and Hoechst33342 co-staining, NPY1R staining was first performed on live cells on ice as above, then cells were fixed with 4% PFA, and permeabilized in 0.1% Triton-X. Finally, Hoechst33342 (10 μg/ml) was added for 20 minutes, and cells were analyzed on the flow cytometer.

### Biochemical Assays

#### Immunoblotting

Cells were lysed in RIPA buffer (50 mM Tris-HCl, pH 7.5, 150 mM NaCl, 2 mM EDTA, 1% NP-40, 0.5% Na deoxycholate, and 0.1% SDS) with protease and phosphatase inhibitors (40 μg/ml PMSF, 20 mM NaF, 1 mM Na_3_VO_4_, 10 mM β-glycerophosphate, 10 mM sodium pyrophosphate, 2 μg/ml antipain, 2 μg/ml pepstatin A, 20 μg/ml leupeptin, and 20 μg/ml aprotinin). Lysates were cleared by centrifugation at 15,000 g for 10 minutes at 4°C, and supernatants were collected. Protein concentrations were quantified by Bradford assay (Thermo Scientific), and 10-30 μg total cellular protein were resolved on SDS-PAGE and transferred onto Immobilon-FL PVDF membranes (Millipore). Bound antibodies were detected with IRDye infrared secondary antibodies using the Odyssey Infrared Imaging System (LI-COR Biosciences). Primary antibodies used for immunoblots included: pHER2 Y1221/1222 (2249; Cell Signaling Technology), c-ErbB2 (OP15L; Millipore), pEGFR Y1068 (2234, Cell Signaling Technology), EGFR (2232, Cell Signaling Technology), pERBB3 Y1289 (4791; Cell Signaling Technology), ERBB3 (sc-285, Santa Cruz Biotechnology), pAKT S473 (4060, Cell Signaling Technology), pPRAS40 T246 (13175; Cell Signaling Technology), p-p70 S6 Kinase T389 (9205; Cell Signaling Technology), p-p44/42 (9101L; Cell Signaling Technology), ERK2 (C-18; Santa Cruz Biotechnology Inc.), p-GSK3β S9 (9336; Cell Signaling Technology), pTSC2 S939 (ab52962; Abcam), pS6 S240/244 (2215; Cell Signaling Technology). pFoxO1 (Thr24)/FoxO3a (Thr32) (9464; Cell Signaling Technology), SGK3 (8156; Cell Signaling Technology), pNDRG1 (T346) (5482; Cell Signaling Technology), p-AKT substrate (9614L, Cell Signaling Technology), TSC2 (3990; Cell Signaling Technology).

#### Kinase Assays

To enrich for dephosphorylated TSC2 to use as a substrate for *in vitro* kinase assays, pcDNA3-*wtTSC2* was transfected into 293T cells, and after 24 hours, transfected cells were treated with 1 μM BKM 120 for 1 hour before lysis in kinase assay lysis buffer (50 mM Tris-HCl, pH 7.5, 150 mM NaCl, 1% NP-40, 1 mM EDTA, 1 mM EGTA, 40 μg/ml PMSF, 20 mM NaF, 1 mM Na_3_VO_4_, 10 mM β-glycerophosphate, 10 mM sodium pyrophosphate, 2 μg/ml antipain, 2 μg/ml pepstatin A, 20 μg/ml leupeptin, 20 μg/ml aprotinin, 0.1% (v/v) 2-mercaptoethanol). FLAG-tagged TSC2 was immunoprecipitated with ANTI-FLAG^®^ agarose beads (Sigma) for 2 hours at 4°C. Beads were subsequently washed once with high-salt kinase assay wash buffer (50 mM Tris-HCl, pH7.5, 500 mM NaCl, 1% NP-40, 1 mM EDTA, 1 mM EGTA, 40 μg/ml PMSF, 20 mM NaF, 1 mM Na_3_VO_4_, 10 mM β-glycerophosphate, 10 mM sodium pyrophosphate, 2 μg/ml antipain, 2 μg/ml pepstatin A, 20 μg/ml leupeptin, 20 μg/ml aprotinin, 0.1% (v/v) 2-mercaptoethanol), once with kinase assay lysis buffer, and twice with kinase assay buffer (50 mM Tris-HCl pH 7.5, 10 mM MgCl_2_, 0.1 mM EGTA). Kinase reactions contained recombinant SGK3 (100 ng), 100 μM ATP, 0.1% (vol/vol) 2-mercaptoethanol, 20 mM NaF, 1 mM Na_3_VO_4_, 10 mM β-glycerophosphate, and 10 mM sodium pyrophosphate. Reactions were allowed to proceed for 30 minutes at 30°C, and then were terminated by boiling in SDS-PAGE sample buffer. Proteins from these assays were resolved with 6-10% SDS-PAGE.

To monitor SGK3 phosphorylation of TSC2 *in vivo*, 293T cells on 60 mm dishes were co-transfected with 1.5 μg GST-SGK3 and 1.5 μg FLAG-tagged TSC2 (wildtype TSC2, TSC2-SATA, TSC2-5A or TSC2-6A). Cells were allowed to recover in regular growth media for 24 hours, and then were lysed in kinase assay lysis buffer. TSC2 was immunoprecipitated with ANTI-FLAG^®^ beads and subjected to immunoblotting with pAKT substrate antibody. Whole cell lysates and immunoprecipitated TSC2 were resolved by 6-15% SDS-PAGE.

### Mass Spectrometry

Scaled-up kinase reaction mixtures were resolved by SDS-PAGE. Gels were stained with Coomassie Brilliant Blue R-250 Staining Solution (BioRad), and a band corresponding to FLAG-TSC2 was excised and submitted to the Perlmutter Cancer Center Proteomics shared resource at NYU School of Medicine (New York, NY, USA). Bands were cut into 1 mm^3^ pieces and de-stained for 15 minutes in a 1:1 (v/v) solution of methanol/100mM ammonium bicarbonate. The buffer was exchanged, and the samples were de-stained for another 15 minutes. This process was repeated for another 3 cycles. Gel pieces were then dehydrated by washing in acetonitrile, dried in a SpeedVac for 20 minutes, reduced in 100μl of 0.02M dithiothreitol (Sigma)/pH 7.5 for one hour at 57 ⁰C, and alkylated with 100μl of 0.05M iodoacetamide (Sigma) for 45 minutes at room temperature in the dark. Gel pieces were once again dehydrated by washing in acetonitrile, and then dried in a SpeedVac for 30 minutes. Sequencing grade modified trypsin (500 ng, Promega) was added directly to the dried gel pieces, followed by enough 100mM ammonium bicarbonate to cover them. The gel plugs were agitated at room temperature, and digestion was allowed to proceed overnight. Reactions were halted by adding a slurry of R2 50 μm Poros beads (Applied Biosystems) in 5% formic acid/0.2% trifluoroacetic acid (TFA) to each sample at a volume equal to that of the ammonium bicarbonate added for digestion. Samples were allowed to shake at 4⁰C for 120 mins, and the beads were loaded onto C18 ziptips (Millipore) equilibrated with 0.1% TFA by centrifugation for 30s at 6,000 rpm in a microfuge. The beads were then washed with 0.5% acetic acid, and peptides were eluted in 40% acetonitrile in 0.5% acetic acid, followed by 80% acetonitrile in 0.5% acetic acid. The organic solvent was removed by using a SpeedVac, and the sample was reconstituted in 0.5% acetic acid.

An aliquot of each sample was loaded onto an Acclaim PepMap trap column (2 cm x 75 µm) in line with an EASY-Spray analytical column (50 cm x 75 µm ID PepMap C18, 2 μm bead size) by using the auto-sampler of an EASY-nLC 1200 HPLC (Thermo Fisher Scientific) with solvent A (2% acetonitrile in 0.5% acetic acid) and solvent B (80% acetonitrile in 0.5% acetic acid). Peptides were eluted into an Orbitrap QExactive HF-X Mass Spectrometer (Thermo Fisher Scientific) by using the following gradient: 5-35% in 120 min, 36-45% in 10 min, followed by 45-100% in 10 min. High resolution full MS spectra were recorded at a resolution of 45,000, an AGC target of 3e6, a maximum ion time of 45ms, and a scan range from 400 to 2000m/z. Following each full MS scan, parallel reaction monitoring (PRM) scans were acquired for the peptides of interest. MS/MS spectra were collected at a resolution of 15,000, an AGC target of 2e5, maximum ion time of 120ms, one microscan, 2m/z isolation window, fixed first mass of 150 m/z, dynamic exclusion of 30 sec, and Normalized Collision Energy (NCE) of 27.

MS/MS spectra were searched against the Uniprot human reference proteome database, (downloaded 10/2016) using Byonic software(95, 96) (Protein Metrics Inc., San Carlos, CA). The mass tolerance was set to 10 ppm for MS1 and 0.02 Da for MS2 searches. A false discovery rate (FDR) of 1% was applied at the protein level. The Byonic search included fixed modifications of carbamidomethylation on cysteine and variable modifications of oxidation on methionine, deamidation on asparagine and glutamine, and phosphorylation at serine, threonine and tyrosine residues. The spectra of the peptides of interest were verified manually to localize phosphorylation sites. Relative quantification of the peptides was calculated using Byologic software (Protein Metrics Inc., San Carlos, CA), which uses extracted ion chromatogram areas (XIC areas). The XIC is automated by Byologic from the Byonic search results of potentially identified peptides.

### Statistical Analysis

Data are expressed as mean ± SEM unless specified otherwise. All statistical analyses were generated by using GraphPad Prism 5. Significance was assessed by two-way ANOVA, one-way ANOVA, or Student’s t-test, as appropriate. For ANOVAs, the post-hoc analysis used is indicated in the figure legend. A p-value of less than 0.05 was considered significant.

### Data availability

RNAseq and scRNA-seq data have been deposited in Gene Expression Omnibus (GEO: GSE155342 and GEO: GSE156246). All other reagents are available upon request from B.G.N or J.M.

### Code Availability

The bespoke Perl script for the barcode library analysis is available from J.M. upon request. The code for NeoALTTO analysis is available at https://github.com/bhklab/DTP_HER2_BC.

## ACKNOWLEDGEMENTS

We thank the PCC Genome Technology Center (GTC) for expert library preparation and sequencing, the Applied Bioinformatics Laboratories (ABL) for providing bioinformatics support and help with data analysis and interpretation, and the Proteomics Laboratory (PRL) for technical support. GTC, ABL, and PRL are shared resources partially supported by the Cancer Center Support Grant P30CA016087 to the Laura and Isaac Perlmutter Cancer Center. This work also used computing resources at the NYU School of Medicine High Performance Computing (HPC) Facility. We thank Neke Ibeh, Zhibin Lu, and Carl Virtanen (Princess Margaret Cancer Center) for bioinformatic analysis of the NPY1R RNA-seq data. We also thank Dr. Alex Toker (Beth Israel Deaconess Medical Center) for providing the GST-SGK3 plasmid. This work was partially supported by NIH grant CA59152 to B.G.N. and CIHR grant MOP-142375 and support from the Ontario Research Excellence Fund to J.M. During part of this study, B.G.N. was a Canada Research Chair, Tier 1, and J.M. holds a Canada Research in Functional Genomics, Tier 1. A.S. received support from ARO grant W911NF1910243 and NIH grant GM124446. J.S. received support from NIH grant P01CA229086. C.A.C. was supported by Frederick Banting and Charles Best Canada Graduate Scholarship and Doctoral Completion Award from the Department of Medical Biophysics at University of Toronto. C.S. is funded by the Fonds de la Recherche Scientifique (FNRS) and D.V. by the Fondation Julie et Françoise Drion.

## Author Contributions

C.A.C. and B.G.N. conceptualized the study. C.A.C., J.M., J.J., and B.G.N. designed the experiments. C.A.C. performed the majority of the initial experiments. J.J. performed most of the experiments for the revision. A.S.M. and B.H. performed expression analysis of the NeoALTTO trial data. D.V. and C.S. provided data from this trial. S.J., A.S., K.R.B., A.K. and A.T. performed bioinformatics analysis. A.N., J.J. J.M. performed the DNA barcode amplifications and sequencing. A.H. and P.M. helped design and perform genomics analyses. A.D. and B.U. performed the mass spectrometry experiments and analyses. K.H.T. performed the mammary fat-pad injections for the limiting dilution assay. C.A.C. and B.G.N. wrote the manuscript, which all authors helped to edit. J.D., K.K.W. and S.A. provided primary HER2 and TN breast tumor scRNA-seq datasets.

## Supplementary Figure Legends

**Supplementary Fig. S1.**
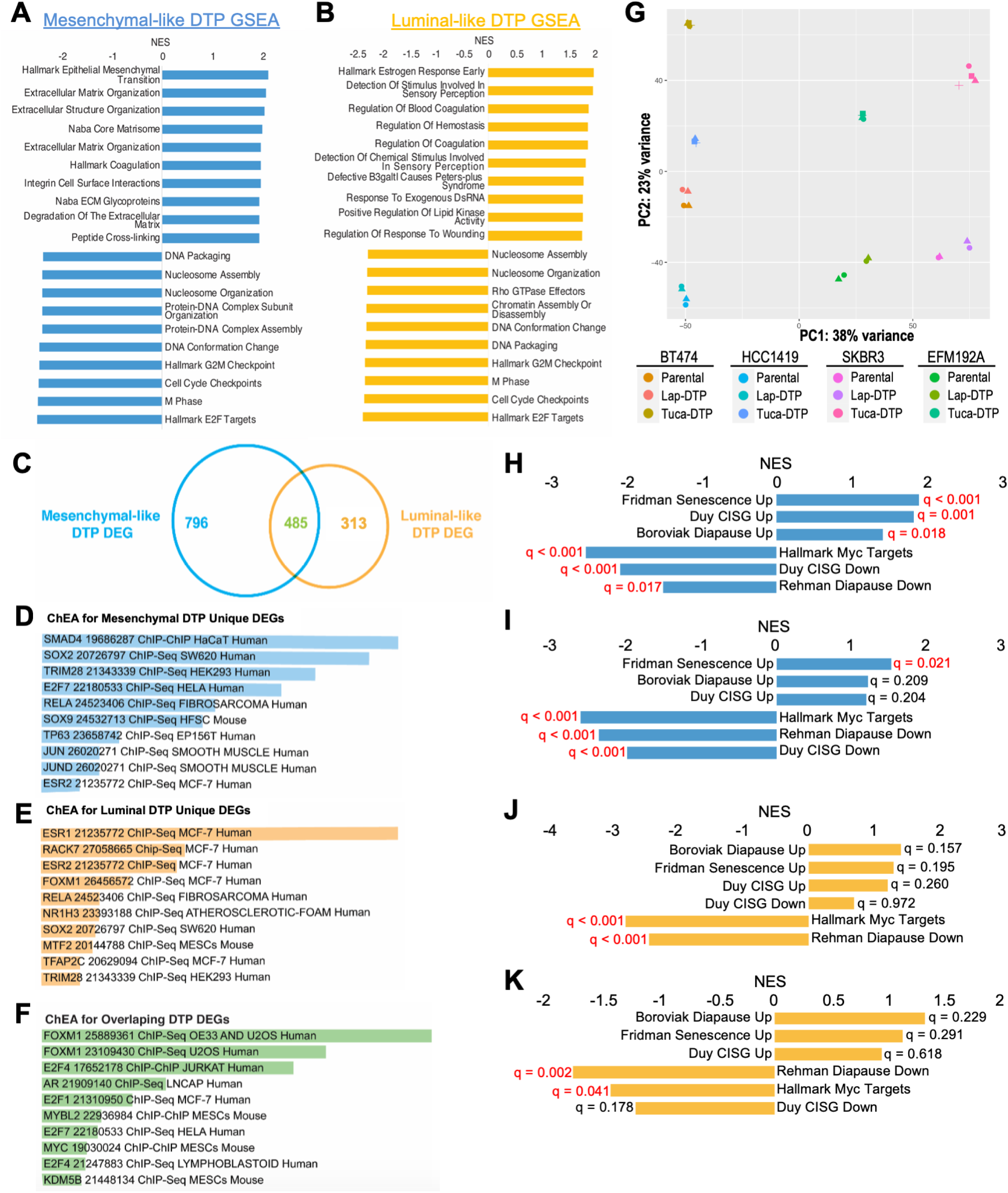
HER2 TKI DTPs exhibit two distinct transcriptional programs, related to Figure 2. **A** and **B,** Top and bottom ten enriched GSEA terms for differentially expressed genes (DEGs) between lapatinib-DTPs and parental cells in the mesenchymal-like (**A**) and luminal-like (**B**) subgroups. **C,** Venn diagram showing overlap of DEGs (compared with cognate parental cells) from mesenchymal-like and luminal-like DTPs. The non-overlapping genes are termed “mesenchymal DTP unique DEGs” (796 genes) and “luminal DTP unique DEGs” (313 genes). **D-F,** TF binding site enrichment, as predicted by ChEA, is displayed for the mesenchymal DTP unique DEGs **(D)**, luminal DTP unique DEGs **(E)**, and overlapping DEGs **(F)**, respectively. Bars indicate ChEA combined score. **G**, Principal component analysis (PCA) of parental cells, lapatinib-DTPs and tucatinib-DTPs from BT474, HCC1419, SKBR3, and EFM192A lines. **H-K**, GSEA for the indicated signatures examined in mesenchymal-like lapatinib-DTPs (**H**), mesenchymal-like tucatinib-DTPs (**I**), luminal-like lapatinib-DTPs (**J**), and luminal-like tucatinib-DTPs (**K**)

**Supplementary Fig. S2.**
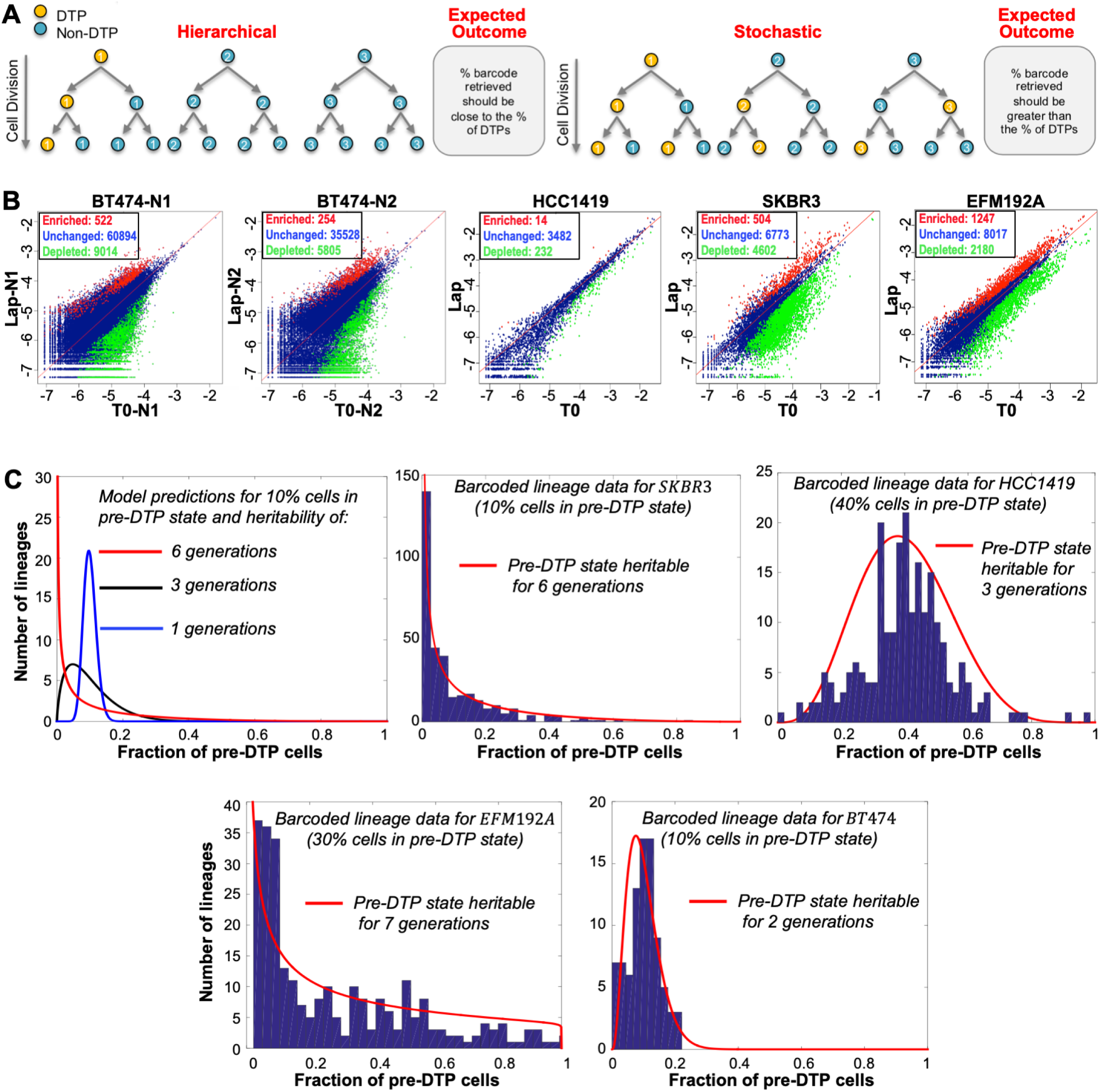
Barcoding experiments indicate stochastic origin of lapatinib DTPs, related to Figure 3. **A,** Schematic shows potential models of DTP ontogeny and expected outcomes of barcoding experiments **B,** Plots show distributions of barcodes expressed as Log(clone frequency) from T0 and 14d lapatinib-treated cells. Red indicates significantly enriched barcodes, green indicates significantly depleted barcodes, and blue shows unchanged barcodes after 14d lapatinib treatment (DTPs). **C,** Model-predicted lineage-to-lineage fluctuations in the fraction of pre-DTPs for different transient heritability of the pre-DTP state. Longer heritability drives enhanced fluctuations (Top left). Distribution of pre-DTPs as obtained from barcoding lineage data from the four cell lines. The corresponding model fits (solid red lines) reveal the pre-DTP state heritability for each cell line.

**Supplementary Fig. S3.**
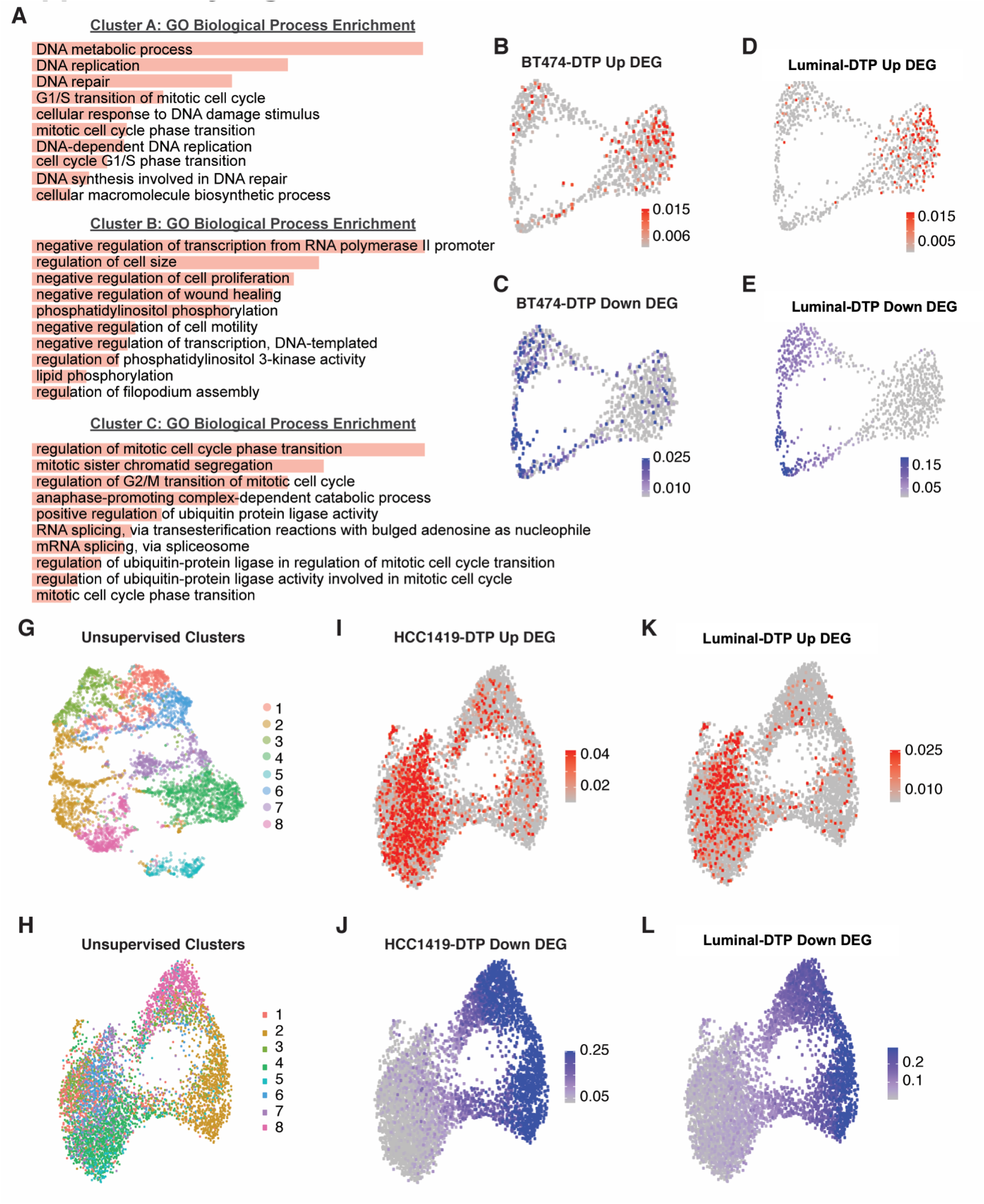
Single-cell RNA-sequencing reveals sub-population enriched for lapatinib DTP genes in untreated HER2+ breast cancer cells, related to Figure 3. **A,** GO Biological Process enrichment for unsupervised clusters of untreated BT474 cells. **B-E,** UMAP of supervised clustering of untreated BT474 cells using cell cycle signature genes defined by Xue *et al*. Cells are colored by BT474-DTP Up (**B**) or Down (**C**) DEG expression or by luminal-DTP Up (**D**) or Down (**E**) DEG expression. Scale shows the signature expression score for each. **G,** UMAP showing the results of unsupervised clustering of untreated HCC1419 cells. Each unsupervised cluster is represented by a number and color, as indicated. **H,** UMAP showing supervised clustering of untreated HCC1419 cells with cell cycle signature genes defined by Xue *et al.*, colored by the unsupervised clusters in **G**. **I-L,** UMAP showing supervised clustering of untreated HCC1419 with Xue *et al.* cell cycle genes, with cells colored by HCC1419-DTP Up (**I**) or Down (**J**) DEG expression or luminal-like DTP Up (**K**) or Down (**L**) DEG expression. Scale shows the signature expression score for each.

**Supplementary Fig. S4.**
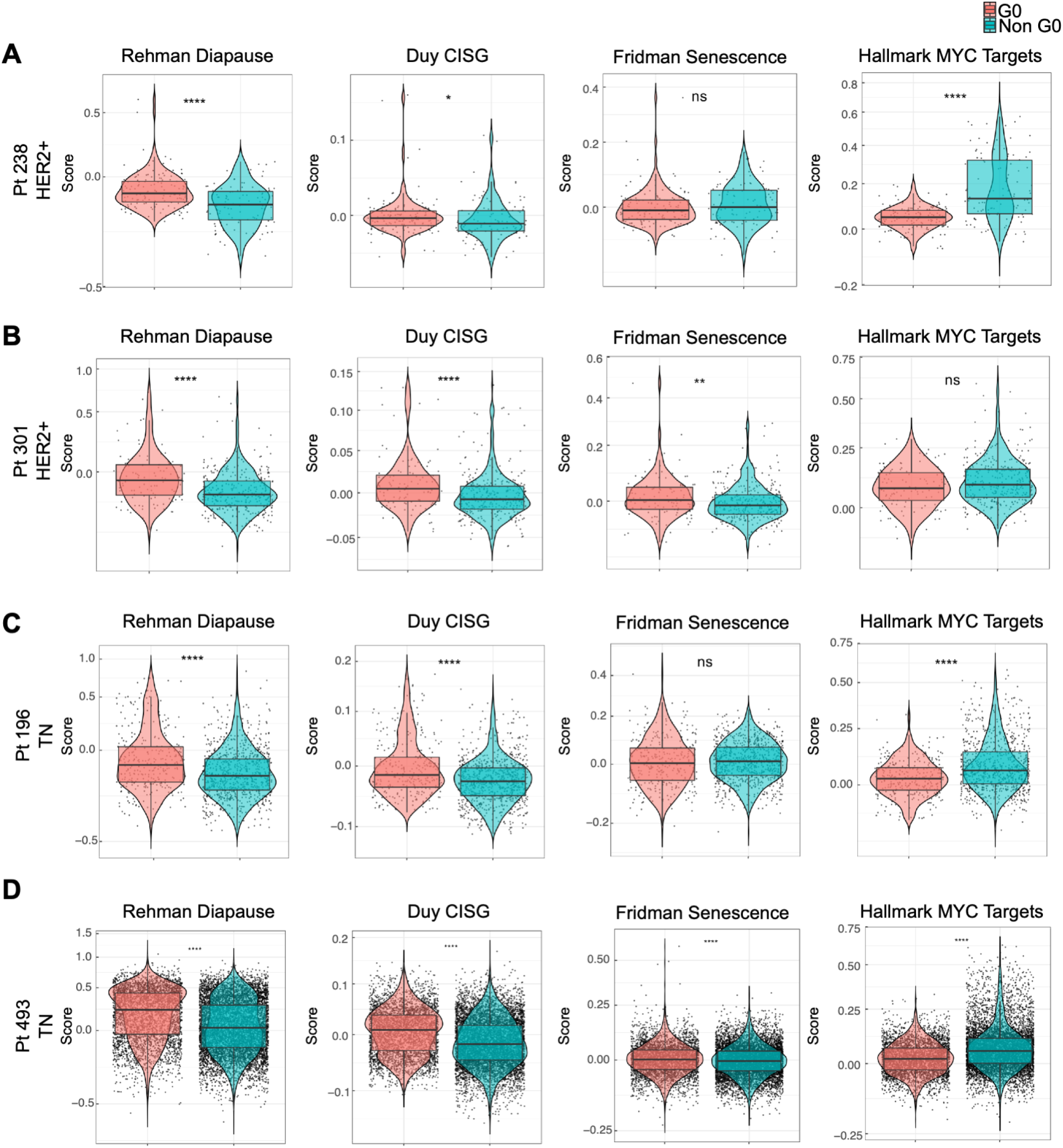
G_0_ cells from treatment-naive primary breast tumors are enriched for diapause genes, chemotherapy-induced stress, and senescence genes, related to Figure 4. Signature scores for Rehman_Diapause_UP, Duy_CISG_UP, Fridman_Senescence_UP, and Hallmark_of MYC_targets in HER2+ tumors from Pt238 (**A**) and Pt301 (**B**), and from TNBC tumors from Pt196 (**C**) and Pt493 (**D**).

**Supplementary Fig. S5.**
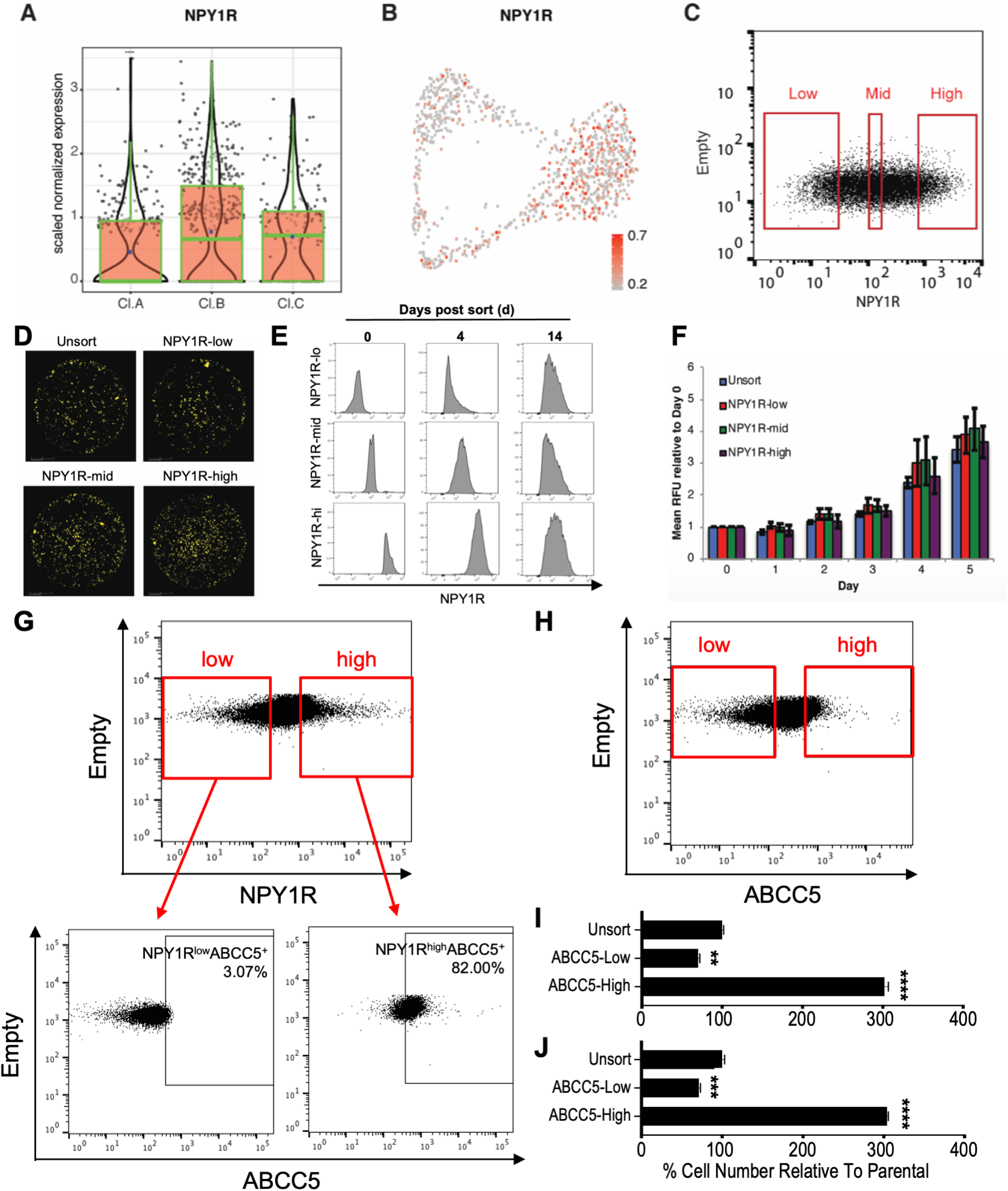
Enrichment of pre-DTPs from untreated BT474 cells by FACS for NPY1R and ABCC5, related to Figure 5. **A,** *NPY1R* expression in unsupervised clusters from untreated BT474 cells. **B,** UMAP showing supervised clustering of untreated BT474 single cells by their cell cycle gene expression with cells colored by *NPY1R* expression level. **C,** Plot showing NPY1R levels in untreated BT474 cells and gates used to define NPY1R^hi^, NPY1R^mid^, and NPY1R^lo^ fractions. **D,** FACS-isolated cells from the indicated NPY1R fractions were treated with lapatinib for 14 days and then cultured in drug-free media for 14 days. Representative Incucyte images at the experimental endpoint are displayed, and a color mask is applied to show residual cells. **E,** NPY1R^hi^, NPY1R^mid^, and NPY1R^lo^ BT474 cells were isolated by FACS and cultured under standard conditions. NPY1R expression was assessed by flow cytometry post-sort (Day 0) and at Days 4 and 14 of culture. **F,** FACS-isolated NPY1R cells were cultured under standard conditions, and viable cell number was quantified by Alamar Blue assay at the indicated times. Alamar Blue readings (RFU) were compared to the Day 0 values for each sample**. G,** Plot showing NPY1R levels in untreated BT474 cells and gates used to define NPY1R^hi^ and NPY1R^lo^ fractions (upper panel). ABCC5 expression is shown for NPY1R^hi^ (lower right) and NPY1R^lo^ (lower left) cells. **H-J,** ABCC5^hi^ and ABCC5^lo^ BT474 cells were isolated by FACS (**H**) and treated with lapatinib (**I**) or tucatinib (**J**) for 14 days. Cells were counted and compared to unsorted control cells. Mean ± SEM from three independent experiments is shown. Significance was assessed by one-way ANOVA with Tukey multiple comparisons test (**p<0.01, ***p<0.001, ****p<0.0001).

**Supplementary Fig. S6.**
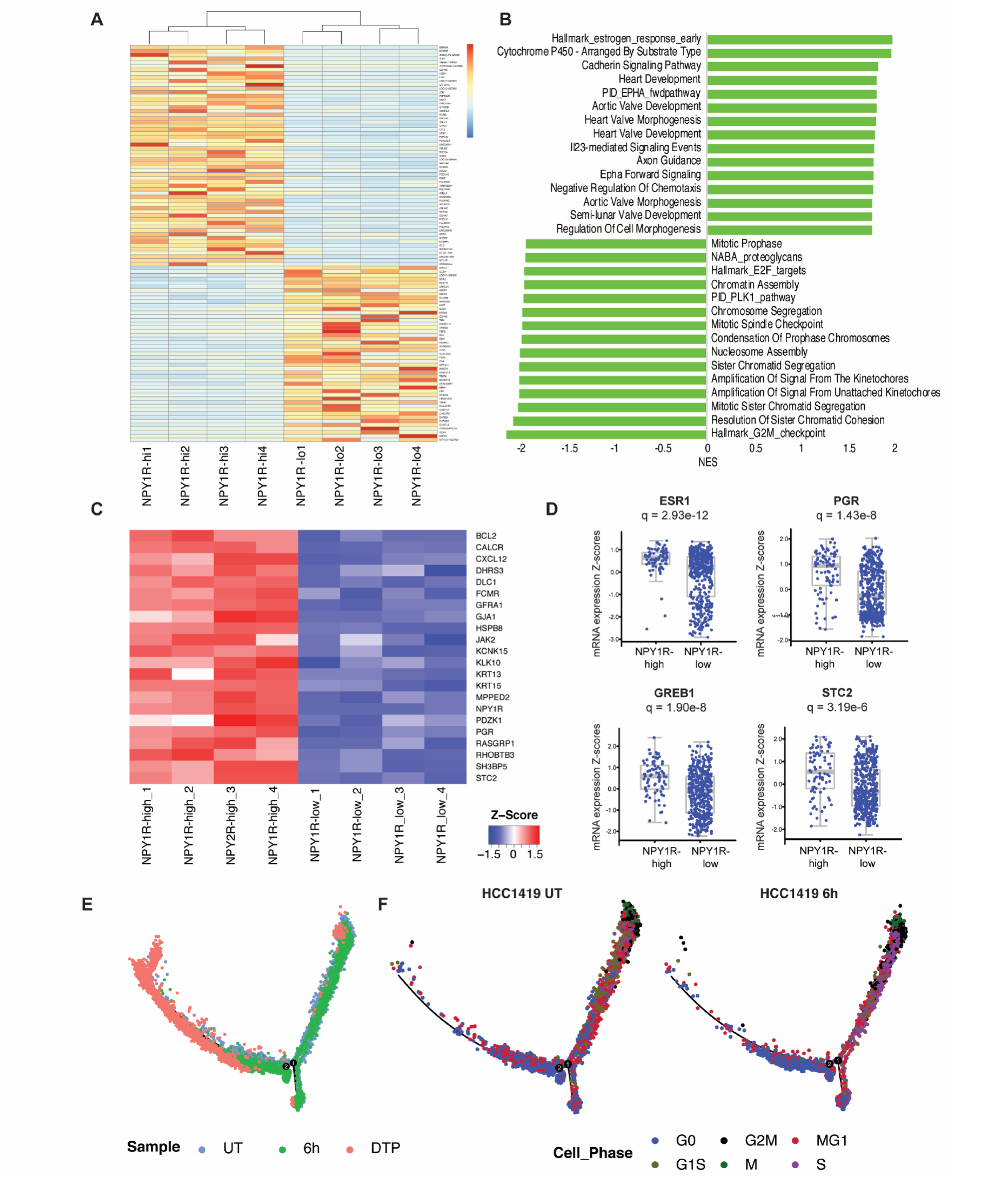
NPY1R^hi^ cells have enhanced basal estrogen receptor (ER) activity and HCC1419 G_0_ cells are on trajectory to become DTPs, related to Figure 5. **A,** Heatmap showing DEGs from NPY1R^hi^, compared with NPY1R^lo^, cells. Scale represents z-score. **B,** Top ten positively and negatively enriched GSEA terms, ranked by normalized enrichment score (NES), for DEGs in FACS-isolated NPY1R^hi^ vs. NPY1R^lo^ BT474 cells. **C,** Heat map showing ER targets enriched in NPY1R^hi^ compared with NPY1R^lo^ cells. Scale represents z-score. **D,** Expression of ER targets in NPY1R^hi^ vs. NPY1R^lo^ breast tumors from TCGA. NPY1R^hi^ tumors are defined as those with *NPY1R* expression ≥ 1 SD above the mean; all other tumors are considered NPY1R^lo^. The q value was calculated by the Benjamini-Hochberg procedure. **E,** Pseudotime analysis of untreated and 6h lapatinib-treated HCC1419 cells and HCC1419 DTPs with cells colored by sample. **F,** Respective positions of untreated and 6h lapatinib-treated HCC1419 cells on the pseudotime trajectory, with cells colored by cell cycle phase, as defined by the Xue *et al*.

**Supplementary Fig. S7.**
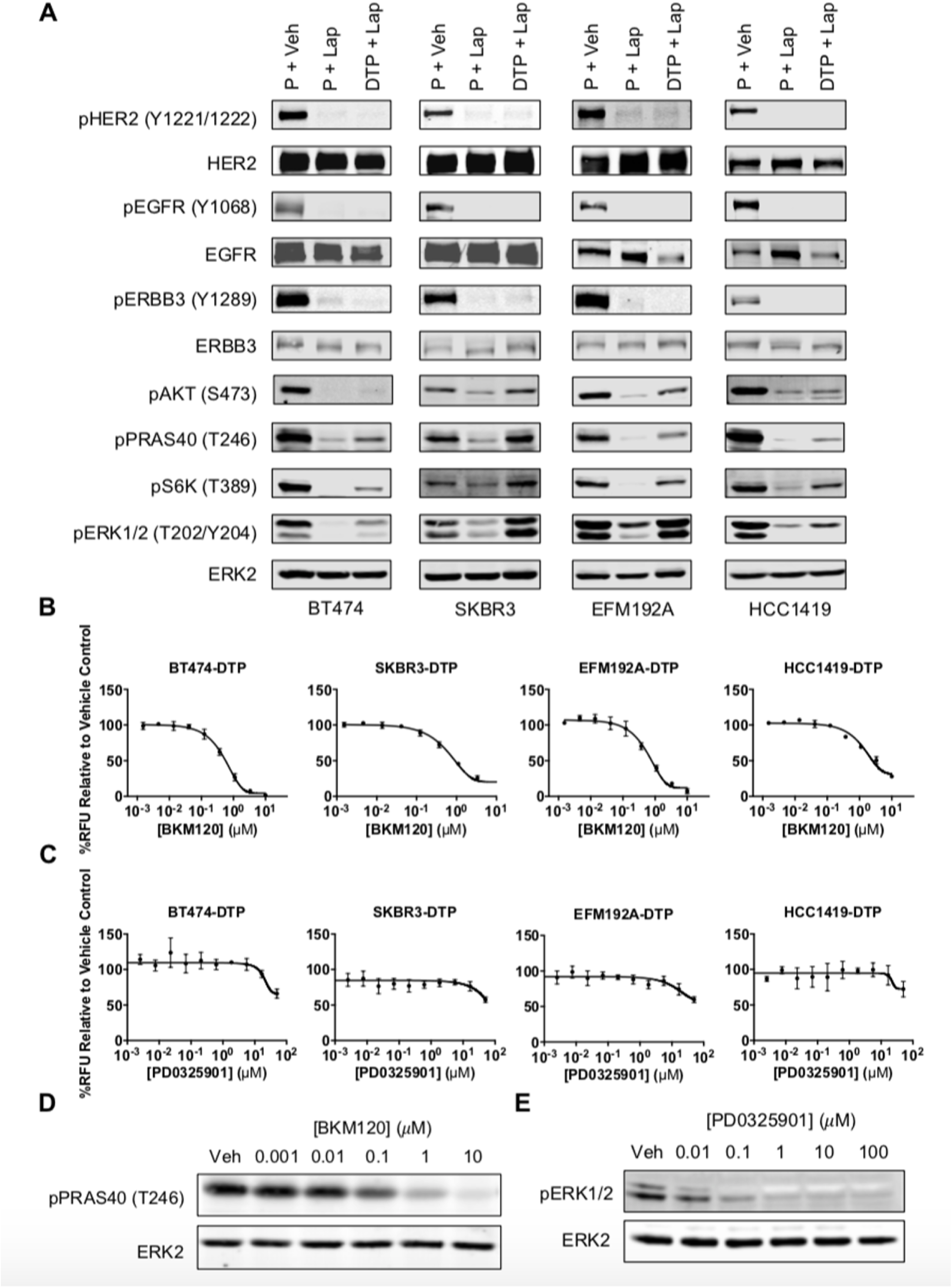
PI3K and ERK/MAPK pathways are re-activated in lapatinib DTPs, but only PI3K pathway is required for lapatinib-DTP survival, related to Figure 6. **A,** Parental cells (P) were treated with vehicle or 2.5 μM lapatinib for one hour (P + lap) and compared with lapatinib-DTPs cultured continuously in 2.5 μM lapatinib. Downstream signaling events were assessed by immunoblotting with the indicated phospho-specific antibodies. ERK2 serves as a loading control. **B** and **C,** Lapatinib-DTPs were treated with the pan-PI3K inhibitor BKM120 (**B**) or the MEK inhibitor PD0325901 (**C**) for 96 hours. Surviving cells were quantified by Alamar Blue viability assay. Mean ± SEM of normalized relative fluorescence units (RFU) from three independent experiments is displayed. **D** and **E,** EFM192A lapatinib-DTPs were treated with the indicated doses of BKM120 (**D**) and PD0325901 (**E**) for one hour, and whole cell lysates were analyzed immunoblotting to assess pathway inhibition. ERK2 serves as a loading control.

**Supplementary Fig. S8.**
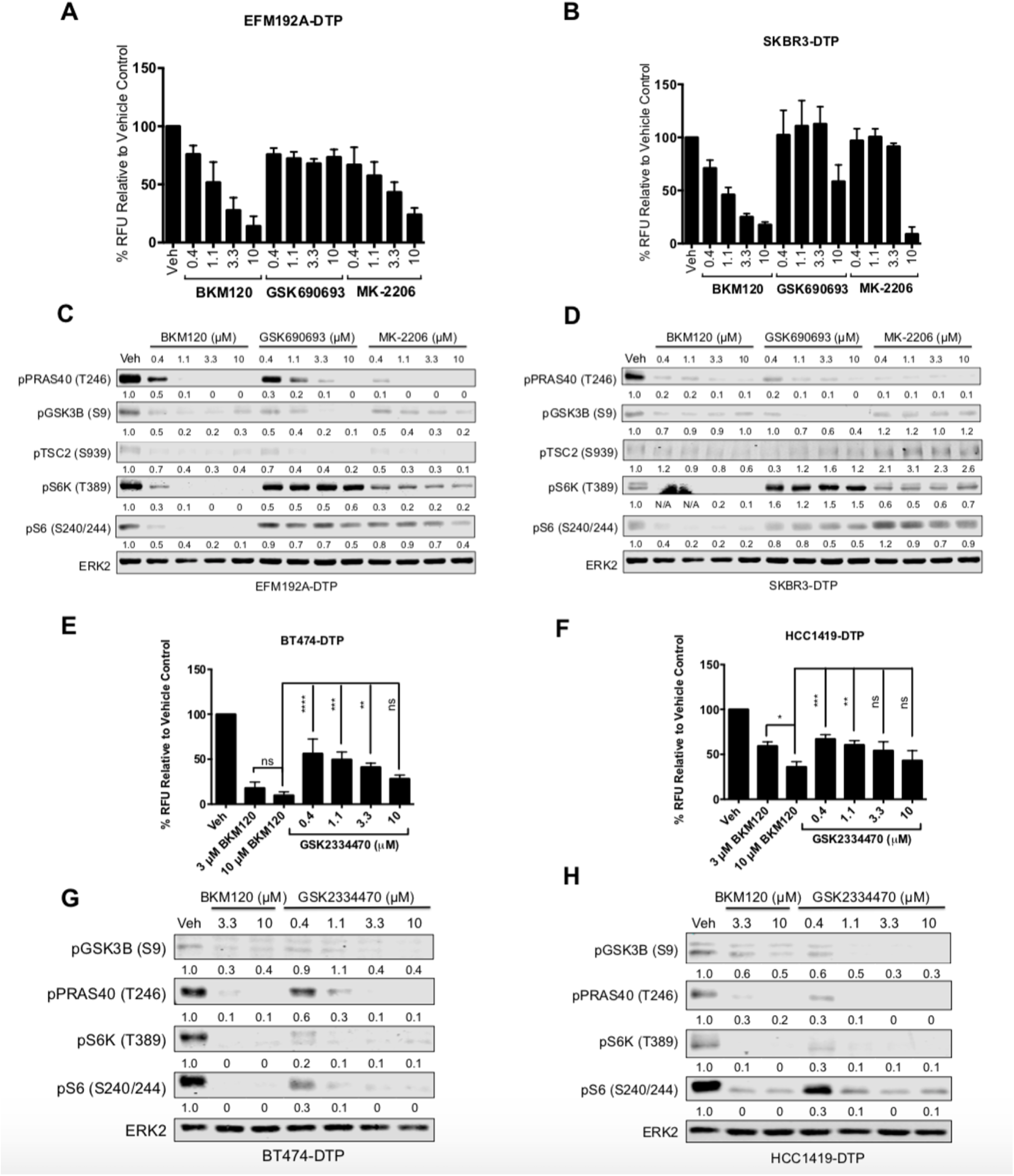
Mesenchymal-like lapatinib-DTPs also are more sensitive to PI3K than to AKT inhibition and mTORC1 activation and survival in luminal-like lapatinib-DTP are dependent on PDK1, related to Figure 6. **A** and **B,** Lapatinib DTPs from EFM192A (**A**) and SKBR3 (**B**) cells were treated with increasing doses of BKM120, GSK690693, or MK-2206 for 96 hours, and surviving cells were quantified by Alamar Blue viability assay. Mean ± SEM of normalized relative fluorescence units (RFU) from three independent experiments is shown. **C** and **D,** EFM192A (**C**) and SKBR3 (**D**) DTPs were treated as indicated for one hour, and whole cell lysates were analyzed by immunoblotting with the indicated antibodies. Numbers under the blots indicate relative intensities compared to the vehicle control. **E** and **F,** Lapatinib-DTPs from BT474 (**E**) and HCC1419 (**F**) cells were treated with indicated doses of BKM120 or the PDK1 inhibitor GSK2334470 for 96 hours, and surviving cells were quantified by Alamar Blue. Mean ± SEM of normalized relative fluorescence units (RFU) from three independent experiments is displayed. (ns, p>0.05, *p<0.05; **p<0.01, ***p<0.001, ****p<0.0001, one-way ANOVA, with Tukey multiple comparisons test). **G** and **H,** Lapatinib-DTPs from BT474 (**G**) and HCC1419 (**H**) cells were treated as indicated for one hour, and whole cell lysates were subjected to immunoblotting. Numbers under the blots indicate relative intensities compared to vehicle control.

**Supplementary Fig. S9.**
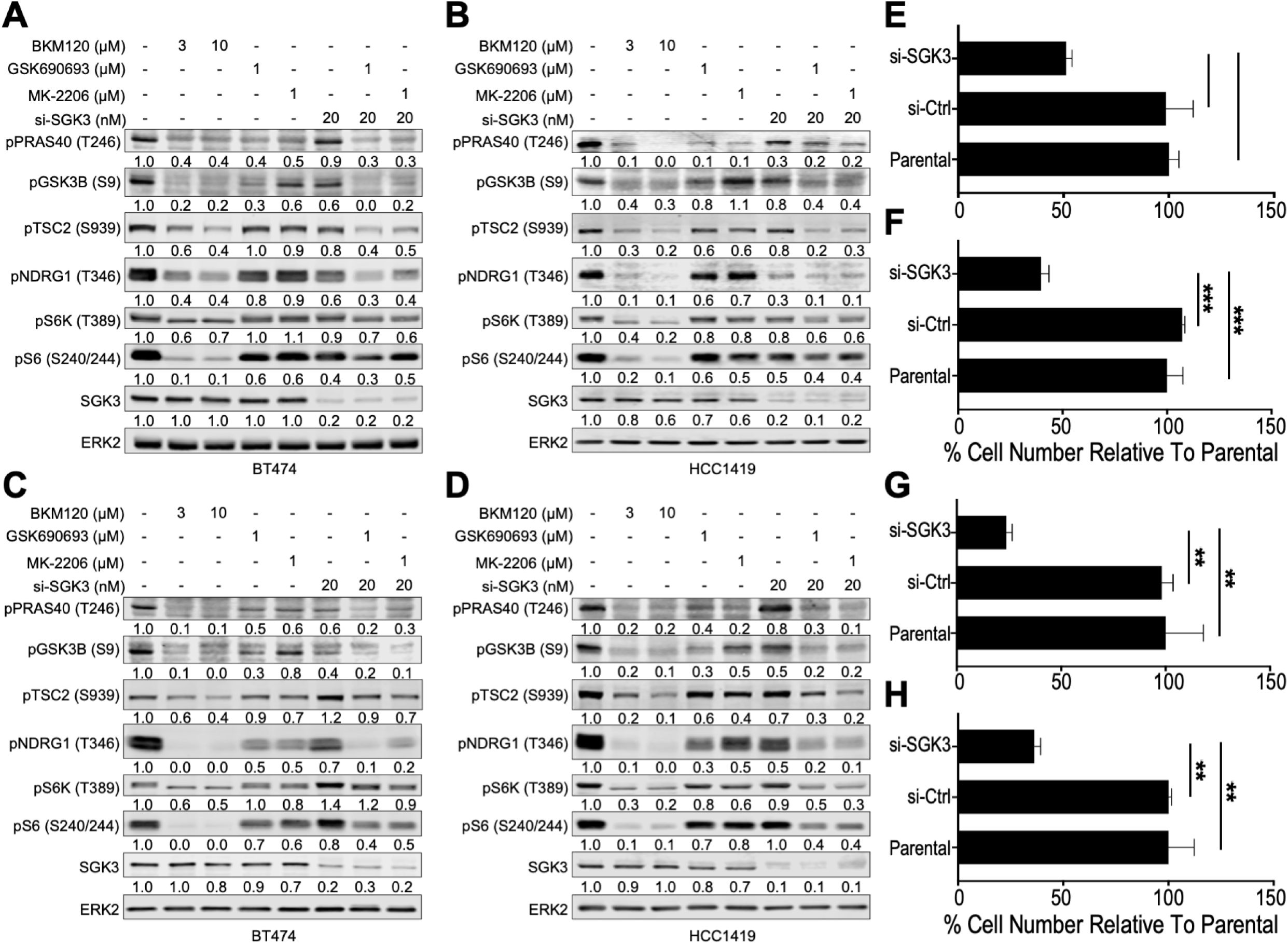
SGK3 knockdown plus AKT inhibition ablates AKT-independent TSC2 phosphorylation and mTORC1 activation and reduces survival of cells in response to HER2 TKIs. **A** and **B,** *SGK3* was depleted by using siRNA (si-SGK3) in BT474 (**A**) and HCC1419 (**B**) for 72 hours before treatment with lapatinib for another 72 hours. Cells were then treated with 3 μM or 10 μM BKM120, 1 μM GSK690693, or 1 μM MK-2206 as indicated for one hour, and whole cell lysates were subjected to immunoblot analysis with the indicated antibodies. Numbers under the blots indicate relative intensity compared to the vehicle control. Representative blots from one of two independent experiments are displayed. **C** and **D,** Same as A and B except cells were treated with tucatinib. Representative blots from one of two independent experiments are displayed. **E**-**H,** Parental BT474 (**E, G**) and HCC1419 (**F, H**) cells or BT474 and HCC1419 cells transfected with si-Ctrl or si-SGK3 for 72 hours were treated with lapatinib (**E** and **F**) or tucatinib (**G** and **H**) for 14 days. Cells were counted and compared to parental control. Mean ± SEM from three independent experiments is shown. Significance was assessed by one-way ANOVA with Tukey multiple comparisons test (*p<0.05, **p< 0.01, ***p<0.001).

